# Optical Control of Translation with a Puromycin-Photoswitch

**DOI:** 10.1101/2022.07.12.499823

**Authors:** Tongil Ko, Mauricio Oliveira, Jessica M. Alapin, Johannes Morstein, Eric Klann, Dirk Trauner

## Abstract

Translation is an elementary cellular process that involves a large number of factors interacting in a concerted fashion with the ribosome. Numerous natural products have emerged that interfere with ribosomal function, such as puromycin, which mimics an aminoacyl tRNA and causes premature chain termination. Here, we introduce a photoswitchable version of puromycin that, in effect, puts translation under optical control. Our compound, termed **puroswitch**, features a diazocine that allows for reversible and nearly quantitative isomerization and pharmacological modulation. Its synthesis involves a new photoswitchable amino acid building block. **Puroswitch** shows little activity in the dark and becomes substantially more active and cytotoxic, in a graded fashion, upon irradiation with various wavelengths of visible light. *In vitro* translation assays confirm that **puroswitch** inhibits translation with a mechanism similar to that of puromycin itself. Once incorporated into nascent proteins, **puroswitch**, reacts with standard puromycin antibodies, which allows for tracking *de novo* protein synthesis using western blots and immunohistochemistry. As a cell-permeable small molecule, **puroswitch** can be used for nascent proteome profiling in a variety of cell types, including primary mouse neurons. We envision **puroswitch** as a useful biochemical tool for the optical control of translation and for monitoring newly synthesized proteins in defined locations and at precise time points.

## INTRODUCTION

Translation is an essential process in cellular growth, plasticity, and differentiation that is particularly active in cancer progression, viral pathogenesis, and memory formation.^1–5^ To study translation, many naturally occurring molecules have been employed that inhibit or stall it, such as cycloheximide, anisomycin, or puromycin. Puromycin structurally mimics the 3’ end of an aminoacylated tRNA (aa-tRNA), featuring an amide linkage between its amino acid and nucleoside moieties, instead of the more labile ester linkage of aa-tRNAs.^6,7^ As such, it exhibits a unique mechanism of action that involves entry into the ribosomal A site and covalent trapping of the nascent polypeptide chain from the P-site. After its incorporation, puromycin cannot be further cleaved by incoming aa-tRNAs, leading to stalling and subsequent premature chain release of the puromycilated peptide.^8,9^

Puromycin has been extensively used in molecular biology and various assays that involve puromycylation have been reported.^10^ In SUrface SEnsing of Translation (SUnSET), antibody-coupled detection of puromycilated peptides on the cell surface is used to study translation fluctuations.^11^ Others include the subcellular localization of translating ribosomes with the ribopuromycilation method (PRM), where puromycilated nascent chains are immobilized on ribosomes by the use of chain elongation inhibitors.^12^ Structural analogs of puromycin have been developed to further broaden the scope of its applications (Figure 1A). For example, O-propargyl-puromycin (OPP) features an alkyne group that allows for subsequent enrichment or visualization of puromycilated proteins through copper-catalyzed “click” reaction with an azide conjugated biotin or fluorophore.^13^

**Figure 1.**
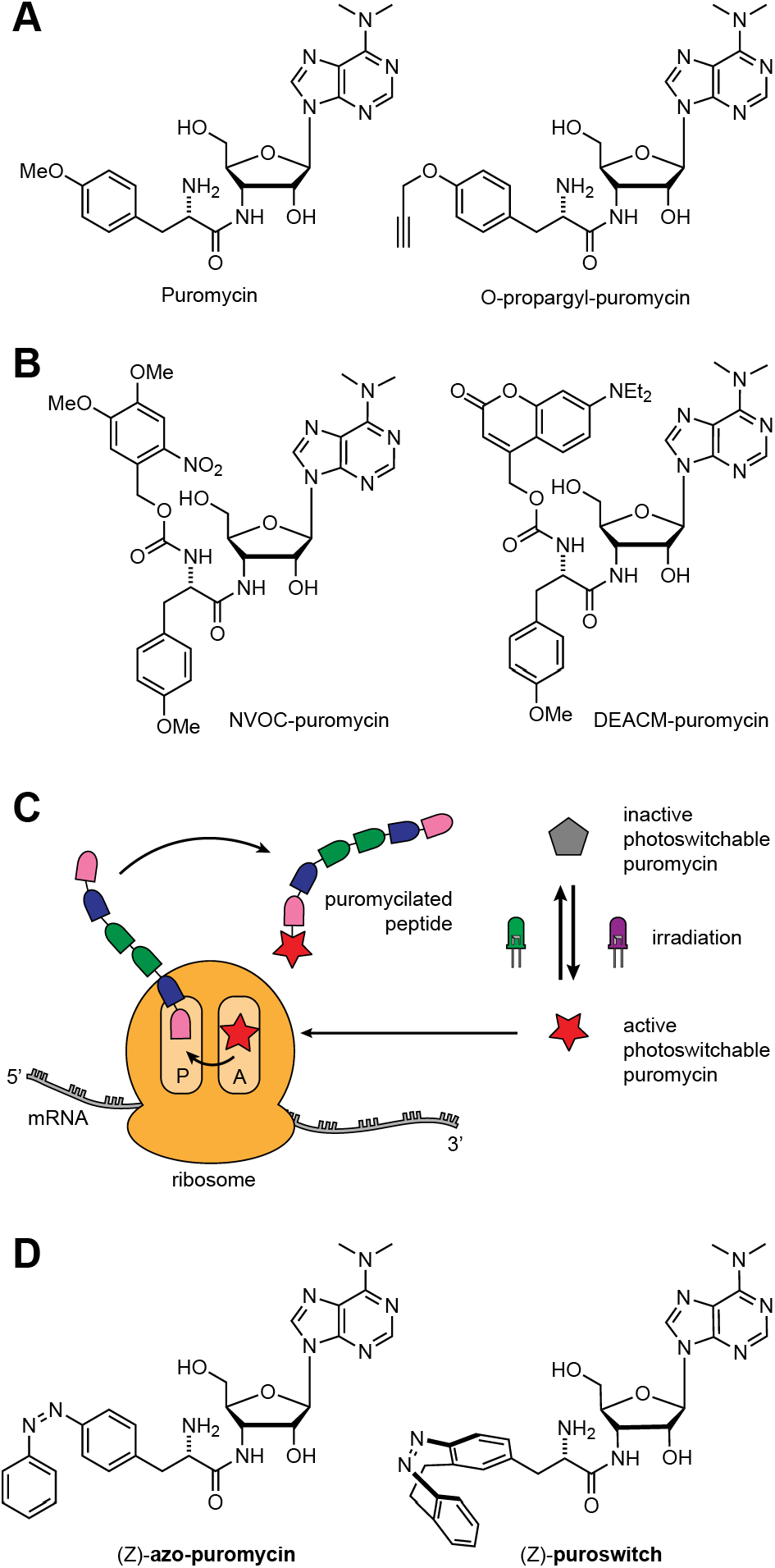
A) Puromycin and functionalized analog O-propargyl-puromycin. B) Photocaged analogs of puromycin. C) Proposed mode of action of photoswitchable puromycin analogs. D) Photoswitchable puromycin analogs synthesized.

In order to allow for spatiotemporal control of the application of puromycin, photocaged versions of the compound have been introduced by the Schwalbe and Schuman groups (Figure 1B). In these compounds, the amino acid position is appended with the photolabile groups O-Nitroveratryloxycarbonyl (NVOC) or 7-Diethylamino-4-methylcoumarin (DEACM).^14,15^ Upon irradiation with UV-A light, puromycin is released. However, photocaging approaches are inherently limited by the irreversible nature of the process where the photocage group, once cleaved, cannot be re-attached. This constraint prevents the spatiotemporal interrogation of processes that occur in specific time intervals as the compound would remain active after release and also diffuse to non-irradiated areas.^16^ Recently, the Schwalbe group developed a once-reversible caged puromycin analog with two photoresponsive groups that could activate and deactivate the compound with two different wavelengths of light.^17^ However, the use of UV-light (365 nm) for the uncaging of these compounds is not ideal for *in vivo* applications as it can have low tissue penetration and prolonged exposure can be damaging to cell cultures and model organisms, making the development of red-shifted tool compounds a desirable goal.^18,19^

Here we describe the development of a photoswitchable puromycin analog that allows for fully reversible and tunable translation inhibition. The compound, which we named **puroswitch**, can be photoisomerized with two wavelengths of visible light, which results in a large difference in biological activity. This feature makes our photoswitchable compound suitable for biological applications where one seeks to understand the role of protein synthesis with temporal and spatial control.

## RESULTS

### Design, synthesis, and photophysical characterization of photoswitchable puromycin analogs

Previously published structure-activity relationship studies showed that the amino acid side chain of puromycin can be modulated to a certain degree without losing activity.^20,21^ In addition, photoswitchable amino acids have been developed for the optical control of peptides and proteins.^22,23^ We chose azobenzene as our photoswitch as it is known for its fatigue resistance, large and predictable geometrical changes, and well-characterized photothermal properties.^24,25^ We thought to “azo-extend” the 4-position of puromycin’s amino acid side chain by substituting the methoxy group with an phenyl diazene moiety, yielding **azo-puromycin** (Fig. 1D).^26^ Intriguingly, this very molecule had already been reported by the Sisido group as spectroscopic standard.^27^ To the best of our knowledge, however, it was never evaluated for its biological activity, especially in a light-dependent fashion. Synthesis of the azobenzene analog was achieved through a HATU promoted coupling of the known Fmoc-protected azobenzene amino acid **1** with the commercially available puromycin aminonucleoside **2**, followed by protecting group removal (Fig. 2A). The photoswitching and thermal relaxation properties of the compound are shown in the Supporting Information (Fig. S2). The optimal wavelength for switching to the *cis* isomer was 370 nm and rapid isomerization back to the *trans* form could be achieved with wavelengths >450 nm.

**Figure 2.**
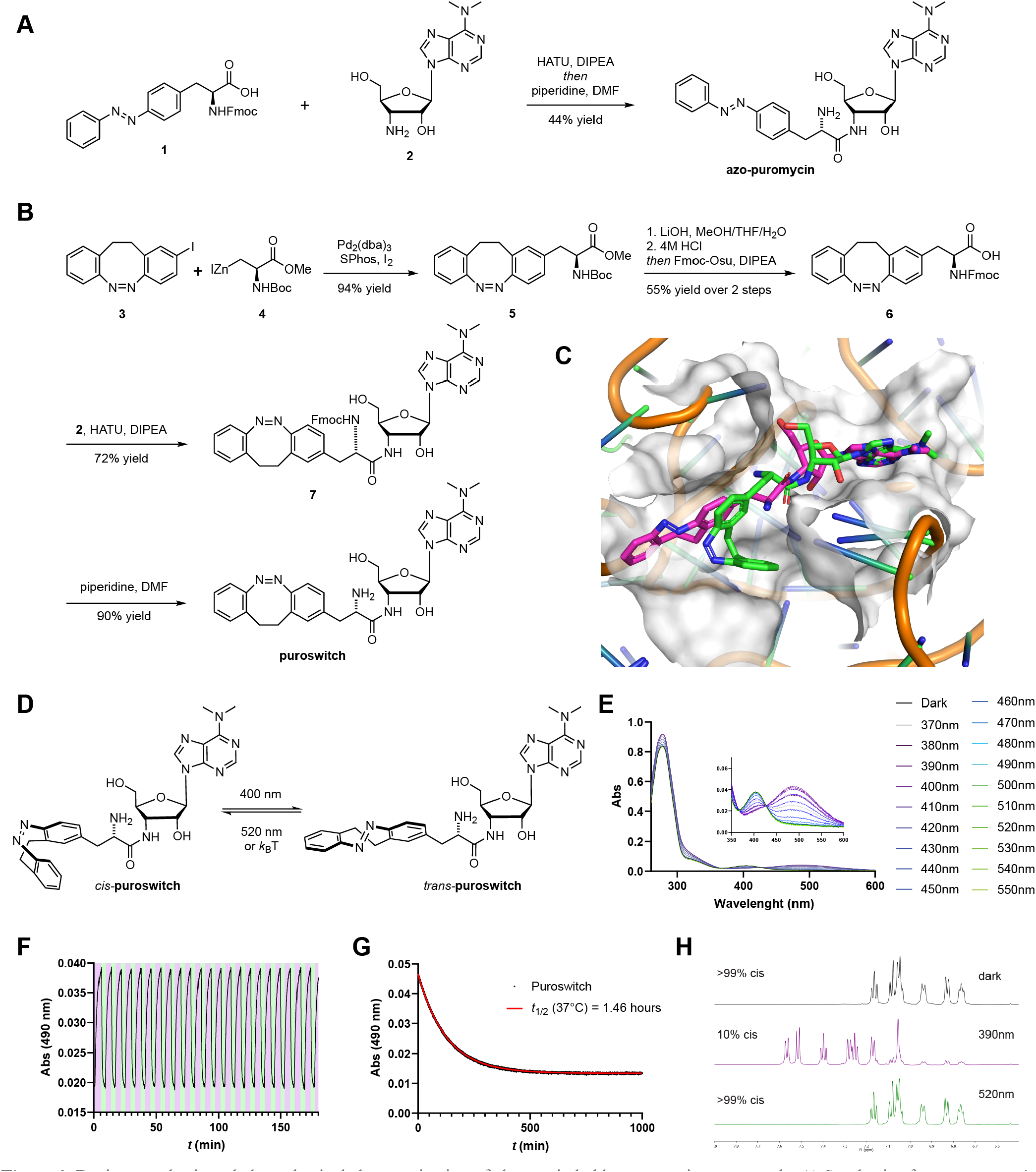
Design, synthesis and photophysical characterization of photoswitchable puromycin compounds. A) Synthesis of **azo-puromycin**. B) Synthesis of **puroswitch**. C) Docking of **puroswitch** in the A site of the ribosome. Best pose from the *cis* (green) and *trans* (purple) **puroswitch** isomers are shown. Model is derived from the structure of CC-puromycin bound to the A site of the 50S ribosomal subunit (PDB: 1Q82). D) Isomerization of *cis*-**puroswitch** to *trans*-**puroswitch**. E) UV-Vis spectra of **puroswitch** (50 μM) in the dark and at different photostationary states in DMSO at r. t. F) Reversible *trans* → *cis* isomerization of **puroswitch** (50 μM) at 390 (purple):520 (green) nm irradiation in DMSO at r.t. G) Thermal relaxation of **puroswitch** (50 μM) at 37°C in DMSO H) ^1^H-NMR spectra of **puroswitch** with different PSS’ in DMSO at r.t.

To assess the light-dependent biological activity of **azo-puromycin**, we tested its effect on mammalian cells. Puromycin has a well-known cytotoxic and growth inhibitory effect in both prokaryotic and eukaryotic cells, allowing us to test its light-dependent activity in a cell viability assay using our ‘Cell DISCO system’ for pulsed irradiation.^28,29^ Human embryonic kidney (HEK) 293t cells were treated with increasing concentrations of puromycin and its analog and were either irradiated with 390 nm for 72 hours at 100 ms every 10s or incubated in the dark. As expected, puromycin showed a similar level of cytotoxicity in both light and dark-adapted conditions (Fig. 3A). **Azo-puromycin** was assayed in similar conditions but with 370 nm light irradiation and exhibited cellular toxicity but was 2-fold less potent than puromycin. Unfortunately, we did not observe light-dependent differences in the toxicity of azo-puromycin (Fig. 3B). This indicated that both photoisomers, despite their different configurations, were active inhibitors of protein synthesis. Moreover, computational docking of both isomers showed that despite the geometric changes in the azobenzene amino acid sidechain, both isomers occupied the puromycin binding pocket of the ribosome in a similar fashion (Fig. S3) Thus, we decided to not further pursue the biological evaluation of this compound.

**Figure 3.**
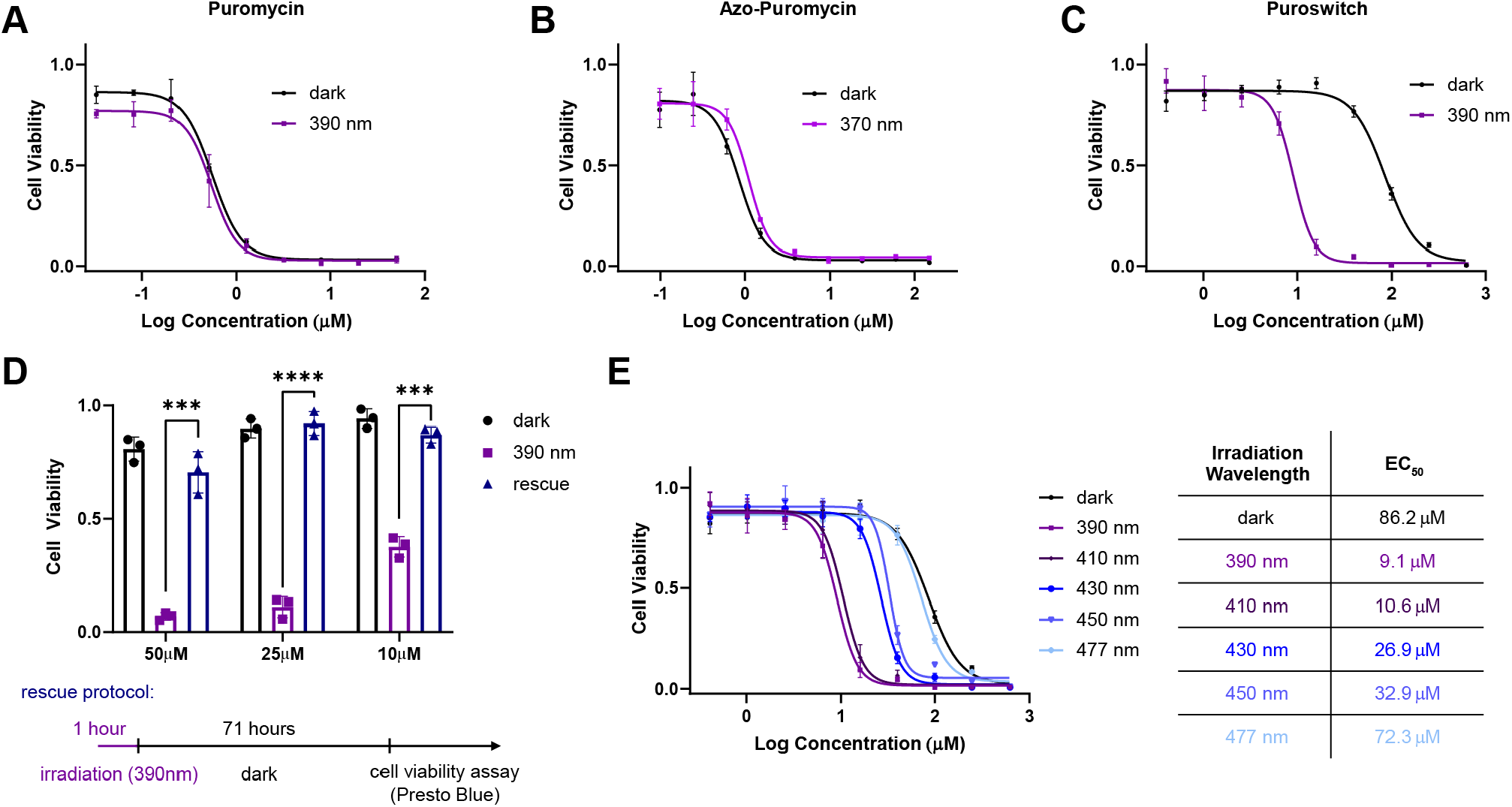
Light-dependent viability of HEK293t cells. A-C) Viability curves for cells treated with different compounds for 72 hours with pulse irradiation (100 ms every 10s) or dark incubation. D) Viability of cells incubated with **puroswitch** under dark, 390 nm irradiation, or rescue protocol. Error bars represent mean values ± SD. Unpaired t-test were performed (****, p<0.0001; ***, p<0.001). E) Viability curves for cells incubated with **puroswitch** under pulse irradiation (100 ms every 10s) at different wavelengths of light.

Inspired by recent success with diazocines in photo-pharmacology, we next synthesized and evaluated a puromycin derivative that we eventually named “**puroswitch**” (Fig. 1D).^30,31^ Diazocines are emerging diaryl diazene photoswitches that, unlike azobenzenes, are more thermodynamically stable in their *cis* configuration. They can be switched back and forth with visible wavelengths of light around 390 nm and 520 nm and reach good photostationary states of more than 90% of the *trans* isomer and almost quantitative amounts of the *cis* isomer.^32^

A computational docking study of **puroswitch** revealed that the *trans* isomer better occupies a binding pocket in the ribosome, closely resembling the binding of puromycin itself, while the *cis* isomer could not properly inhabit that binding pocket (Fig. 2C; Fig. S3). Additionally, we speculated that the bent geometry of the cis isomer could shield the amine functional group from forming amide bonds with the growing peptide chain and consequently diminishing its effect on translation. Isomerization to the *trans* isomer would better expose the amine and allow for our puromycin analog to be incorporated into the nascent peptide chain.

The synthesis of **puroswitch** began with the mono-iodinated diazocine **3**, which was obtained in three steps through our previously reported route.^30^ Subsequent Negishi-Jackson coupling with iodoalanine derived organozinc iodide **4** afforded diazocine **5** in an excellent yield.^33^ After ester hydrolysis and amide coupling with **2**, deprotection of the Boc group and purification of the resulting amino acid proved to be challenging. Thus, we decided to exchange protecting groups in a one pot reaction by treating the hydrolysis product with 4M HCl, followed by addition of Fmoc-Osu and a base, which gave **6**. Finally, amide coupling with **2** and Fmoc deprotection proceeded smoothly to afford **puroswitch**.

**Puroswitch** was shown to optimally isomerize to the *trans* isomer with 390 nm irradiation, where a PSS of 1:9 *cis*:*trans* could be achieved. Rapid *trans* to *cis* isomerization could be achieved by irradiating with 525 nm light, achieving a PSS of >99:1 *cis*:*trans* (Fig. 2E). When kept in the dark, the *trans* enriched diazocine relaxes to the more thermodynamically stable *cis* form with a thermal half-life of 1.46 hours at 37°C in DMSO (Fig. 2D). This feature is desirable as the compound is relatively bistable but also capable of relaxing to its inactive form in the dark. As usually observed for diazocine photoswitches, many cycles of photochemical isomerization were possible without any noticeable fatigue (Fig. 2C).

### Puroswitch exhibits a large difference in bioactivity and enables chromodosing

The light-dependent biological activity of **puroswitch** in HEK293t cells was assessed using cell viability assays as earlier described. The median effective concentration (EC_50_) was determined to be 86.2 μM in the dark and 9.1 μM when irradiated with 390 nm, resulting in a 10-fold difference in activity (Fig. 3C). These results indicate that the compound becomes more toxic upon irradiation, which matched our hypothesis that the *cis* compound could effectively inhibit translation.

Next, we tested the reversibility of our compound. To this end, we designed a rescue assay where HEK293t cells were treated with **puroswitch** at 50, 25 and 10 μM concentrations and immediately pulse irradiated with 390 nm. After one hour, the samples were incubated in the dark to allow **puroswitch** to slowly relax to its dark-adapted *cis* state over 71 hours. Subsequently, cell viability was measured with Presto Blue™ for both set of samples and normalized to that of DMSO treated cells (Fig. 3D). The results showed that samples exposed to irradiation with 390 nm for 72 hours experienced considerable cytotoxicity, while the samples that were allowed to relax back to the *cis* form after 1 h showed viability similar to those kept in the dark.

Another characteristic property of photoswitches and potential advantage over photocaged alternatives is that the concentration of active species can be titrated with the color of the incident light as the PSS is a function of the wavelength (“chromodosing”).^29,34^ To illustrate this point, we ran a chromodosing experiment where we incubated HEK293t cells with different concentrations of **puroswitch** and illuminated the cells with different wavelengths of light. We found that the EC_50_ of **puroswitch** could be gradually tuned with the color of incident light (Fig. 3E).

### Puroswitch enables optical control of translation

To confirm that the cytotoxic effect of **puroswitch** in cells was caused by translation inhibition, we used a rabbit reticulocyte lysate based *in vitro* translation assay.^35^ In this assay, the translation machinery of the cell is assembled *in vitro* and luciferase mRNA is added to the mixture. The resulting synthesis of the luciferase protein is probed through luminescence. As expected, addition of 10μM of puromycin to the mixture led to an almost negligible luminescence signal (Fig. 4A). Incubation of **puroswitch** at 30μM concentration in the dark afforded equivalent levels of luminesce to that of our DMSO control. Irradiation of **puroswitch** with 390 nm light, however, led to significantly reduced levels of luminesce, similar to those observed for puromycin itself. These experiments validate that our compound acts analogously to puromycin, effectively inhibiting translation when irradiated with violet light.

**Figure 4.**
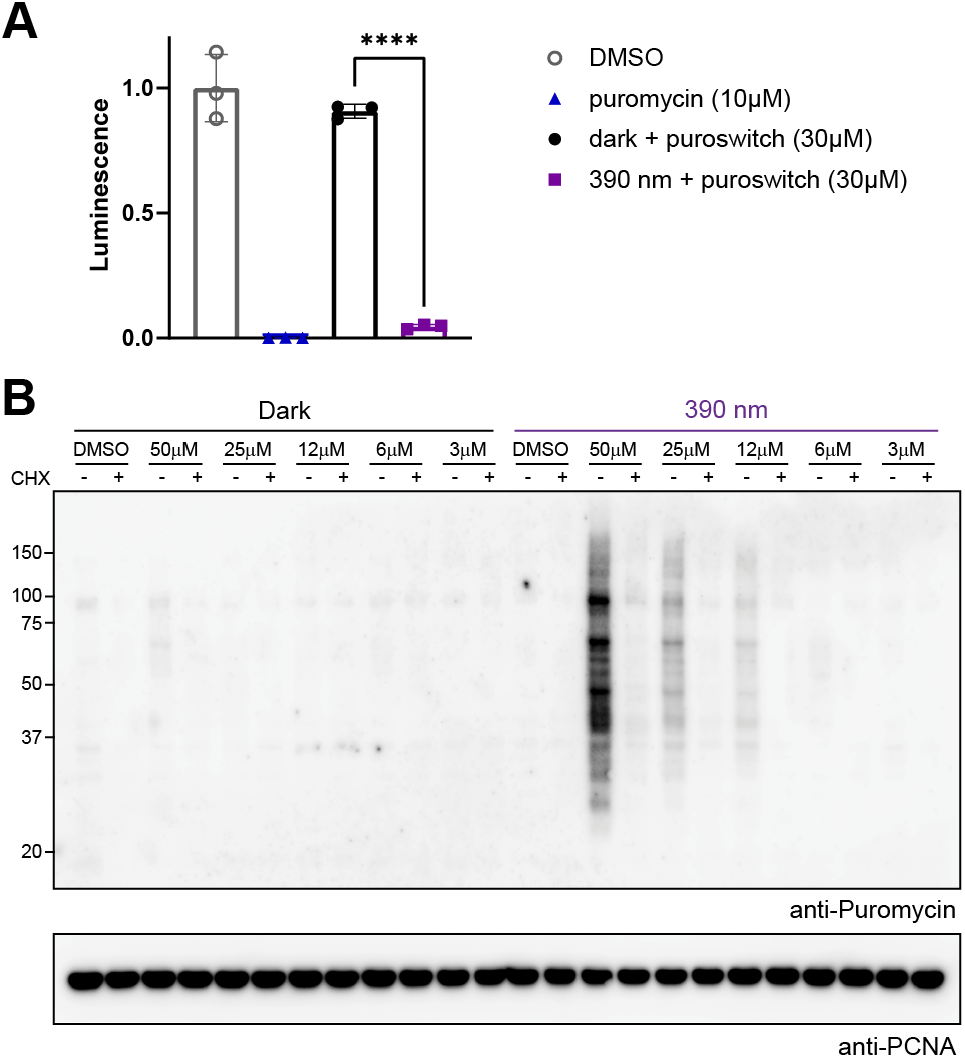
A) Luciferase activity measured for rabbit reticulocyte lysate systems incubated with DMSO, puromycin, or **puroswitch. Puroswitch** samples were irradiated (395 nm) for 2 minutes or left in the dark. Error bars represent mean values ± SD. Unpaired t-test were performed. (****, p<0.0001) B) Time-dependent inhibition of translation. Luciferase activity was measured for samples containing **puroswitch** (30 μM) or DMSO. Samples were irradiated (395 nm) at 29 minutes of incubation for 1 minute. C) Immunoblot analysis after treatment of HEK293t cells with **puroswitch** for 4 hours in the dark (left) or under irradiation (right) at different concentrations. Cycloheximide (CHX) was also added as a control.

### Puroswitch can be used to monitor *de novo* translation of mRNAs

As a consequence of its mechanism of action, which involves incorporation into nascent polypeptide chains, puromycin can be used to monitor *de novo* translation using bioorthogonal chemistry or immunofluorescence.^11,36–38^ Functionalized puromycin analogs that bear handles for biorthogonal chemistry have been used visualization and enrichment through affinity purification.^13,39^ Puromycin antibodies can bind, albeit usually with lower affinity, to analogs containing chemical modifications in the amino acid side-chain.^40^ To test if those antibodies could also bind to **puroswitch**, we treated HEK293t cells with our compound at different concentrations. Western blot analysis using puromycin antibodies showed no signal when the cells were incubated in the dark but a clear concentration dependent puromycilation of proteins in the cells that received 390 nm irradiation (Fig. 4B). To ensure that this process is translation dependent, we pre-treated a control group with cycloheximide (CHX), a translation inhibitor. The fact that we observe significant amounts of puromycilated proteins at relatively high molecular weight points to the fact that *trans* **puroswitch** is considerably less active than puromycin itself, allowing aminoacyl tRNAs to effectively compete until the ribosome encounters a stop signal and pauses. Taken together, these results show that standard puromycin antibodies bind to our compound and confirm that conjugation of **puroswitch** to nascent protein chains can be controlled with light.

### Puroswitch can be incorporated into newly synthesized proteins of primary neurons

Next, we tested the usefulness of **puroswitch** for measuring and controlling mRNA translation in neurons. To assess this, we cultivated mouse primary neurons until maturation and then exposed them to **puroswitch** for 1 h, with or without irradiation with 390 nm light. To confirm that the signal seen was indeed a reliable measure of *de novo* protein synthesis, we ran control experiments in which translation was inhibited with CHX. We found that activation of **puroswitch** with light greatly increased its incorporation into peptides. This effect was significantly attenuated in the presence of CHX, confirming that the signal seen was due to *de novo* protein synthesis (Figure 5). In the dark, we observed incorporation of **puroswitch** at much lower levels. Again, the signal could be completely abrogated by treatment with CHX.

**Figure 5.**
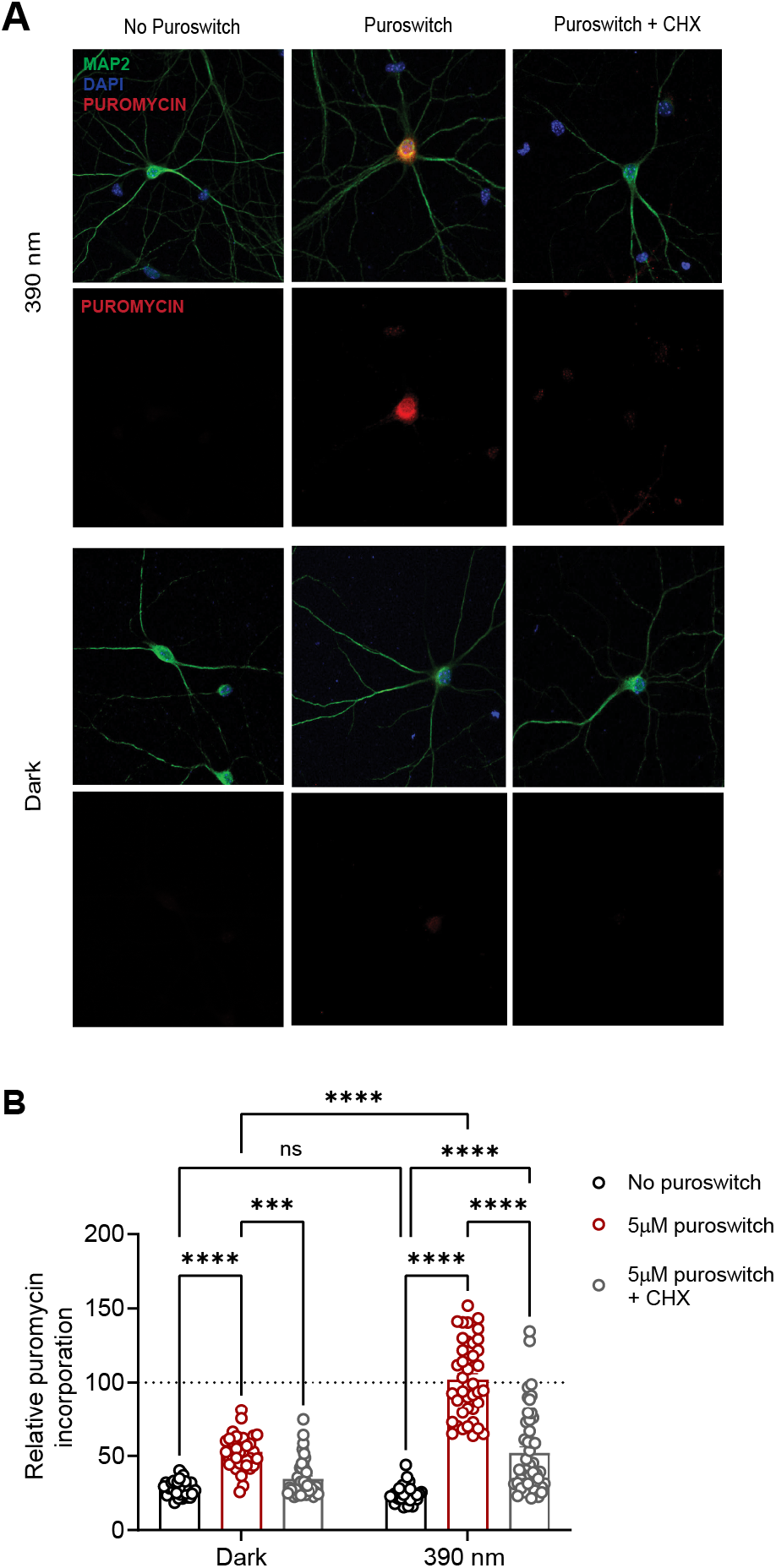
**Puroswitch** incorporation into newly synthesized proteins of rat primary neurons. A) Microscopic immunocytochemistry images of primary neurons stained with anti-MAP2 antibodies (cyan), DAPI (blue), and anti-puromycin antibodies (red). Neurons were incubated with or without **puroswitch** (5 μM) for 1 hour with 390 nm irradiation (100 ms every 10s) or in the dark. Cycloheximide (CHX) was also added as an additional control. B) Relative **puroswitch** incorporation for different compound treatments. Error bars represent mean values ± SEM. Two-way ANOVA with Tukey’s multiple comparisons were performed (****, p<0.0001; ***, p<0.001).

## DISCUSSION

Here, we have introduced an analog of puromycin that contains an a diazocine photoswitch and have evaluated its biological effects in a light dependent fashion. Its synthesis involves the photoswitchable amino acid precursors **5** and **6**, which should be of broad utility to those interested in photoresponsive peptides and proteins. Buildings blocks of this type are easily accessible by combining the methodology for oxidative cyclization of ethylenedianilines with Negishi-Jackson couplings.^30,33^ Diazocine side-chain amino acids are complimentary to classical azobenzene containing ones, first introduced by the Goodman group, which have been used to control various biological functions.^23,41^ The Fmoc protected building block **6**, in particular, should be compatible with solid phase peptide synthesis.

**Puroswitch** builds on and transcends existing approaches to control and monitor translation with caged puromycin derivatives. Photoswitchable probes can be reversibly activated and deactivated and have favorable ON kinetics. **Puroswitch** is stable over many switching cycles and does not generate additional species that can be light-absorbent or toxic. Additionally, diazocine photoswitches have high quantum yields and require low-light intensities in the visible range to isomerize. **Puroswitch** could be effectively isomerized *in vitro* and in cell culture with standard LED’s (<10 mW/cm^2^). Finally, the activity of **puroswitch** can be titrated with the color of incident light. This process, which we term chromodosing, allows for the precise control over the concentration of the active species, a feature that can be difficult to achieve with conventional pharmacology or caged ligands.

**Puroswitch** can be used as a reliable tool for nascent proteome profiling in cells as complex and sensitive as neurons. Neurons are fast-responsive cells, and localized changes in synaptic proteome composition are thought to be necessary for a wide range of processes, such as synapse formation, synaptic pruning, and memory consolidation.^42^ The features of **puroswitch** are well matched with these processes providing a suitable tool to study *de novo* translation in specified locations and at precise time points. We believe that **puroswitch** will empower the development of more complex puromycin-based assays and generate new insights into the time course and location of cellular translation.

## ACKNOWLEDGMENT

T.K. is supported by the NYU MacCracken Fellowship. J.M. thanks the National Cancer Institute (NCI) for a F99/K00 award (K00CA253758). This research was supported by the National Institutes of Health (NIH) grant NS122316 and the NYU Grossmann School of Medicine.

## TOC

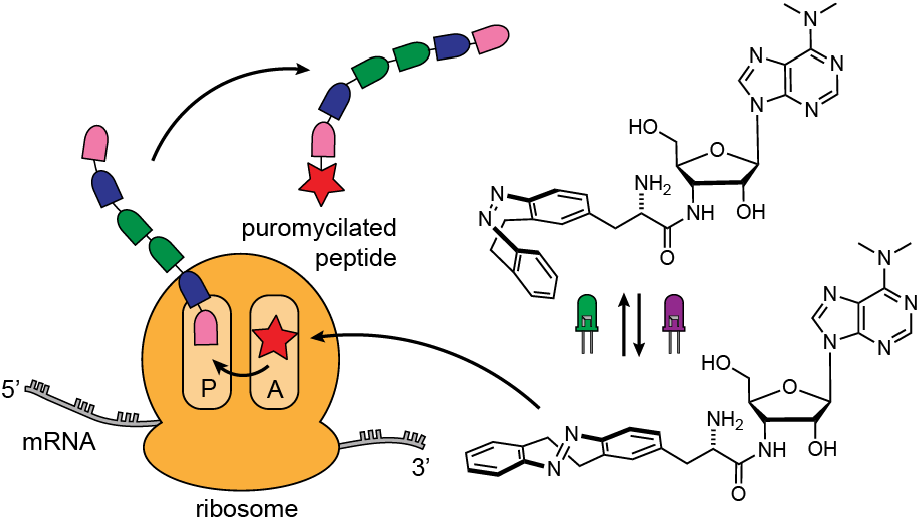

