## Supplementary Information for "Optical Control of Translation with a Puromycin-Photoswitch"

for

### Table of Contents

Supplemental Figures S3

General Information and Protocols S6

Synthetic Procedures and Characterization S11

NMR spectra S22

References S33

#### Supplemental Figures

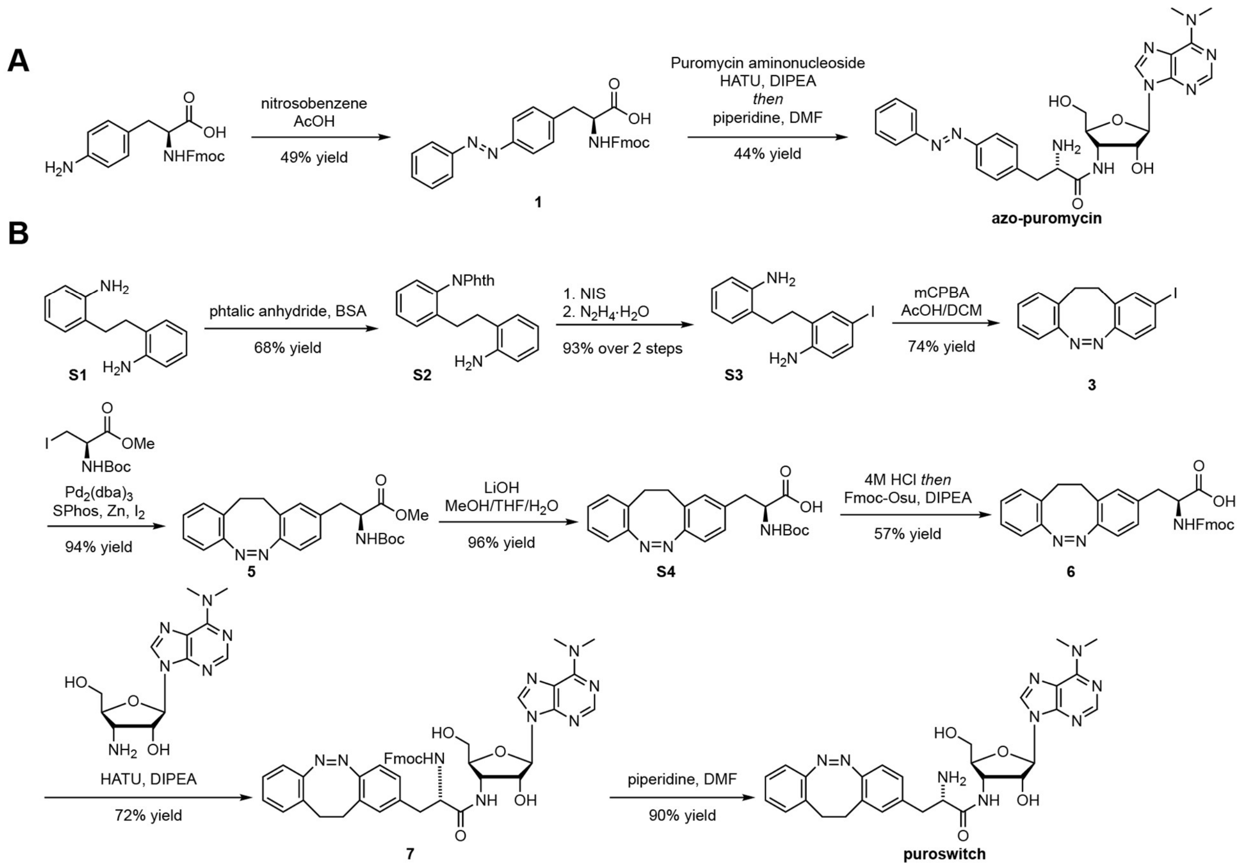


**Figure S1.** Synthesis of **azo-puromycin** (A) and **puroswitch** (B).

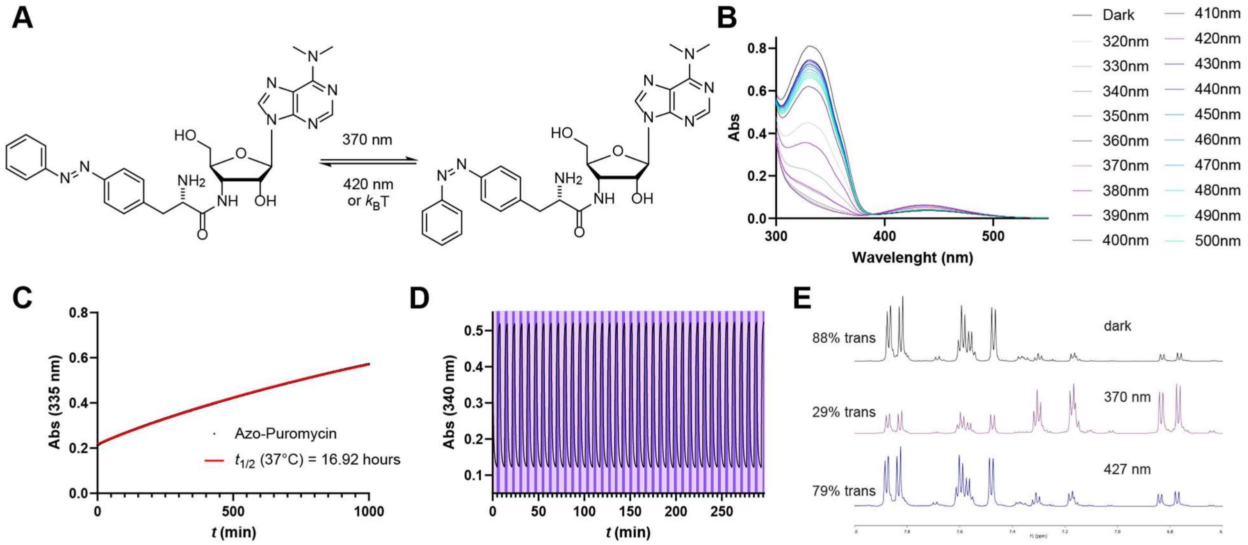


**Figure S2.** Photophysical characterization of **azo-puromycin**. A) Isomerization of *trans*-**azo-puromycin** to *cis*-**azo-puromycin**. B) UV-vis spectra **of azo-puromycin** (50 $\mu$M) in the dark and at different photostationary states in DMSO at r. t. C) Thermal relaxation of **azo-puromycin** (50 $\mu$M) at 37°C in DMSO. D) Reversible *trans*:*cis* isomerization of **azo-puromycin** (50 $\mu$M) at 370 (light purple):420 (purple) nm irradiation in DMSO at r.t. E) $^1$H-NMR spectra of **azo-puromycin** in different PSS in DMSO at r.t.

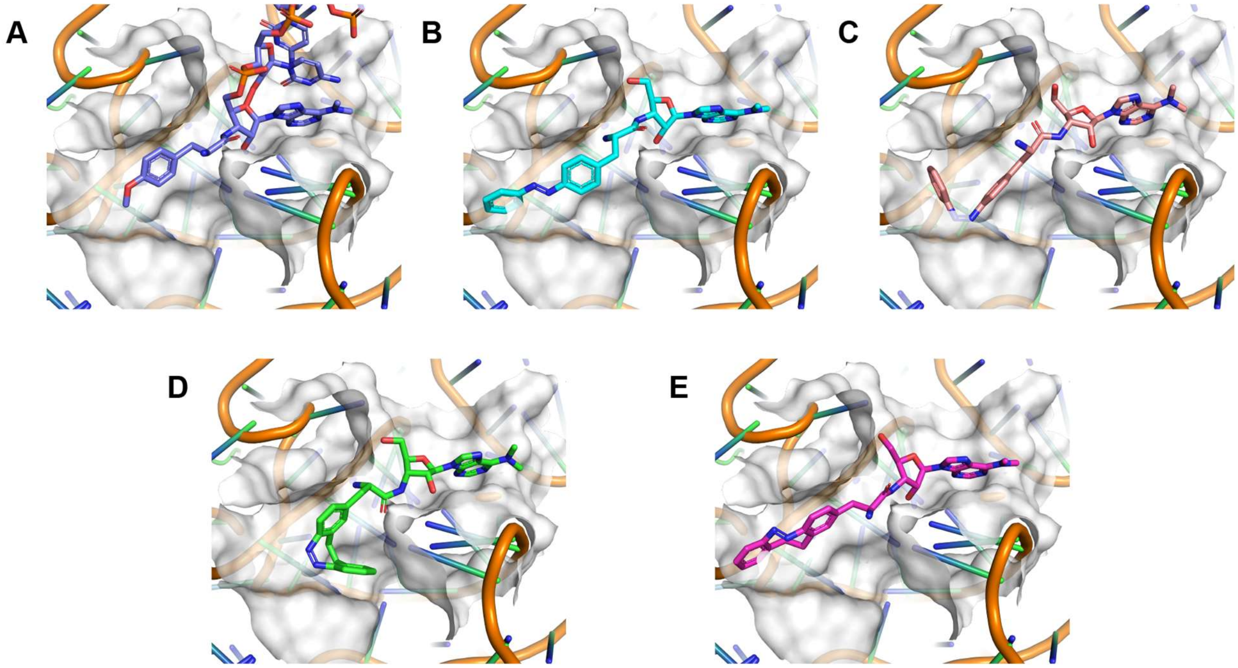


**Figure S3.** A) Crystal structure of CC-puromycin bound to the A site of the 50S ribosomal subunit (PDB: 1Q82). B-E) Docking of photoisomers *trans*-**azo-puromycin**, *cis*-**azo-puromycin**, *cis*-**puroswitch**, and *trans*-**puroswitch** in the A site of the ribosome.

#### General Information and Protocols

##### General experimental conditions

All reactions were conducted with magnetic stirring at room temperature unless otherwise noted. Reactions at elevated temperatures were performed using an oil bath or aluminum block as the heat transfer medium, with the reported temperature corresponding to that of the heat transfer medium.

##### Chemicals

All chemicals were obtained from common vendors and used without further purification unless otherwise noted. With the exception of dimethyl sulfoxide, which was purchased from Oakwood Chemical in standard quality, all solvents were purchased with "certified ACS" or higher quality. Anhydrous solvents were prepared with a solvent purification system by filtration of HPLC grade solvents through alumina or purchased from Acros Organics in AcroSeal™ bottles.

##### Chromatography

Retardation Factors (Rf) were determined by analytical thin-layer chromatography (TLC) performed on pre-coated glass plates from Millipore Sigma (TLC Silica Gel 60 Plates, 250 µm layer thickness, F254 fluorescence indicator), with visualization by exposure to ultraviolet light (254 nm) or by staining with a basic potassium permanganate solution. Column chromatography was performed with silica gel obtained from Millipore Sigma (Geduran® Si 60, 40 – 63 µm). Column chromatography was either performed manually or with an automated chromatography system (Teledyne Isco CombiFlash®).

##### Nuclear magnetic resonance (NMR) spectroscopy

Proton and carbon ($^{1}$H- and $^{13}$C-) NMR spectra were recorded on Bruker Avance III (600/150 MHz, with TCI Cryoprobe™) and Avance III HD (400/100 MHz, with BBFO Cryoprobe™) spectrometers. All spectra were recorded at a temperature of 25 °C in 5 mm tubes in deuterated solvents purchased from Cambridge Isotope Laboratories, Inc. (chloroform-*d* or $CDCl_3$, 99.8% D; dimethyl sulfoxide-$d_6$ or DMSO-$d_6$, 99.9% D). For 1H-NMR spectra chemical shifts (δ) in parts per million (ppm) relative to tetramethylsilane (δ = 0 ppm) are reported using the residual protic solvent ($CHCl_3$ in $CDCl_3$: δ = 7.26 ppm, DMSO-$d_5$ in DMSO-$d_6$: δ = 2.50 ppm) as an internal reference. For $^{13}$C-NMR spectra, chemical shifts in ppm relative to tetramethylsilane (δ = 0 ppm) are reported using the central resonance of the solvent signal (CDCl3: δ = 77.16 ppm, DMSO-$d_6$: δ = 39.52 ppm) as an internal reference. The abbreviations used for multiplicities and descriptors are s = singlet, d = doublet, t = triplet, q = quartet or combinations thereof, m = multiplet and br = broad. NMR spectral data was analyzed with the program MestreNova.

#### Mass spectrometry (MS)

**Liquid Chromatography-Mass Spectometry (LCMS):** Samples were measured on an Agilent Technologies 1260 II Infinity connected to an Agilent Technologies 6120 Quadrupole mass spectrometer with ESI ionization source. Elution was performed using a gradient from 5:95% to 100:0% MeCN:$H_2O$ with 0.1% formic acid over 5 min, if not indicated otherwise. Separated isomer spectra of azobenzenes were obtained by irradiation of the LCMS sample prior to injection.

**High-resolution mass spectra (HRMS):** Spectra were obtained with an Agilent 6224 Accurate Mass time-of-flight (TOF) LC/MS system using either an electrospray ionization (ESI) or atmospheric pressure chemical ionization (APCI) ion source. All reported data refers to positive ionization mode.

#### Computational Docking

Molecules were docked on solved crystal structure of CC-puromycin bound to the A-site of the 50S ribosomal subunit (PDB 1Q82) using Schrodinger software. The crystal structure was truncated by deletion of residues 50Å away from the ligand binding site to minimize computational resources needed. The *cis* and *trans* isomers of **azo-puromycin** and **puroswitch** were prepared using the program LigPrep and the OPLS-3 all-atom force field was applied to all ligands. Docking simulations were performed by Glide with the cubic docking grid being defined as the center of CC-puromycin binding pocket in the ribosome. Constraints were also applied to ensure that the ligands would be docked with the amino acid and nucleotide portions in a similar area to that of CC-puromycin. The best fitting pose was then selected for each ligand based on their GlideScore values.

#### Photophysical characterization

**UV-Vis Spectroscopy**: UV-Vis spectra were recorded on a Varian Cary 60 Scan UV-Vis spectrometer equipped with a Peltier PCB-1500 Thermostat and an 18-cell holder using Brand disposable UV cuvettes (70-850 µL, 10 mm light path) by Brandtech Scientific Inc. Sample preparation and all experiments were performed under red light conditions in a dark room. All UV-Vis measurements were performed with dimethyl sulfoxide (DMSO) as the solvent.

**Wavelength scan:** Light at different wavelengths was provided by an Optoscan Monochromator with an Optosource (75 mW lamp), which was controlled through a program written in Matlab. Irradiation to establish the photostationary state took place from the top through a fiber-optic cable. For each compound a 10 mM stock solution in DMSO was prepared and diluted to a 50 µM concentration prior to the experiment. Spectra with illumination were acquired from 600 to 370 nm in 10 nm steps going from higher to lower wavelengths and illuminating 5 minutes for each wavelength.

**Thermal relaxation:** Compounds were preirradiated with 390 nm light and observing the absorption at 490 nm over 12 hours at 37°C in tightly sealed cuvettes.

#### LED Illumination

**Cell disco:** For illumination of the cells, the cell disco system as previously described in the literature was used.[1] Light-emitting diodes (LEDs) were purchased from Roither Lasertechnik. Pulsed irradiation was performed using 100-ms pulses every 10 s in 96- or 12-well plates, controlled by an Arduino system.

**Sample irradiation:** For cell-free translation experiments, samples were irradiated with UltraFire® UV flashlight (395nm) at specified lengths of time.

#### Cell culture

**Human cell lines:** Human embryonic kidney (HEK) 293t (ATCC, CRL-3216) cell line was purchased from the American Type Culture Collection (ATCC) and cultured in Dulbecco's Modified Eagle Medium (Gibco) with 10% fetal bovine serum and 1% penicillin/streptomycin in a humidified incubator at 37°C with 5% $CO_2$ in air. For the experiments, compounds were serially diluted in cell culture medium without the dye phenol red (Gibco) to reduce the influence of the dye by absorption. Photoswitch stocks and dilutions were strictly kept in the dark and prepared under red light conditions.

**Primary neuron cultures:** Timed pregnant C57/BL6J mice were euthanized at E17 embryonic developmental day. Embryos were recovered in cold Hanks' Balanced Salt Solution (HBSS). Brains were collected and placed in cold HBSS supplemented with glucose. Cortical and hippocampal tissues were dissected from brainstem and meninges were carefully removed with fine tweezers under a low magnification microscope. Brain tissue was cut using fine scissors and placed in 0.25% trypsin (Thermofisher) for 5 min at 37°C. Supernatant was carefully aspirated, tissue was rinsed with warm DMEM + 10% FBS to inactivate any remaining trypsin, and fresh, warm DMEM + 10% FBS was added to the tissue. Brain tissue was mechanically disrupted by pipetting using a fire-polished Pasteur pipette, and then the suspension of cells was separated from remaining tissue using a cell strainer. The total amount of cells was determined using a Neubauer chamber, and cells were plated on coverslips coated with poly-D-lysine + laminin (Sigma-Aldrich). For 18 mm coverslips, 200,000 cells were plated per well. The plates were returned to the incubator (37°C with 5% $CO_2$ in air) for 1 h to allow cell attachment to the glass surface, DMEM (Thermofisher) was aspirated and then replaced with Neurobasal Plus media (Thermofisher) supplemented with B27 plus (Thermofisher), Pen/Strep (Thermofisher) and GlutaMax (Thermofisher). Cells were kept at 37°C with 5% $CO_2$ in air during maturation for 14 days, and regularly fed with fresh supplemented Neurobasal Plus.

#### Presto Blue Cell Viability Assay

The activity of dehydrogenase enzymes in metabolically active cells, as a quantitative measurement for cytotoxicity and proliferation, was determined by fluorescence measurement of the reduction of resazurin to the fluorescent resorufin. The absorbance of resorufin at 570 nm was measured on a FLUOstar Omega microplate reader (BMG Labtech). HEK 293t cells were plated at a density of 10,000 cells in 90 μl of cell culture media and allowed to adhere overnight to a clear 96-well plate. Those cells were then treated with 10 μl of compound stocks at different concentrations in triplicates, using DMSO (1% of final volume) as cosolvent, and incubated for 72 hours. The plates were placed in light-proof boxes and exposed to the lighting conditions specified in the experiment for 72 hours. Next, 10 μl of PrestoBlue™ Cell Viability Reagent (Thermo Fisher, A13261) was added to each well and incubated for 45 to 60 minutes at 37°C. The absorbance at 570 nm was then recorded with a 96-well plate reader. Data was analyzed using GraphPad Prism version 9.11 (GraphPad Software Inc.) and fitted using the sigmoidal dose-response (variable slope) fit. Results represent the mean viability ± SEM relative to the 1% DMSO–treated control.

**Cell Free Translation Assay**

Cellular components for translation from Rabbit Reticulocyte Lysate, Nuclease Treated (Promega, L4960) system and luciferase mRNA (Promega) were assembled in strip polymerase chain reaction (PCR) tubes. Compound stocks were added, using DMSO (1% of final volume) as cosolvent, in triplicates. All reactions had a final volume of 25 μl, contained 500 ng of mRNA and were supplemented with RNasin® Ribonuclease Inhibitor (Promega, N2111) to prevent the degradation of sample mRNA by contaminating RNases. Reactions were immediately irradiated (395 nm for 2 minutes) or left in the dark. The samples were then incubated at 30°C for 90 minutes before 2.5 μl reaction samples were added to 50 μl of Luciferase Assay Reagent (Promega, E1500) in a 96-well plate. Next, the bioluminescence was measured on a FLUOstar Omega microplate reader (BMG Labtech). Data was analyzed using GraphPad Prism version 9.11 (GraphPad Software Inc.). Results represent the mean luminescence ± SEM relative to the 1% DMSO–treated control.

**Puroswitch Incorporation by Western Blotting**

Incorporation of puroswitch in HEK 293t cells was assessed by Western blotting with anti-puromycin antibodies. Cells were platted into 12 well plates and grown to 70-80% confluence in 0.9 ml of cell growth media. Then 100 μl of compound stocks at different concentrations, using DMSO (0.1% of final volume) as cosolvent at different concentrations were added into each respective well. After 4 hours of incubation at different illumination conditions, the media was aspirated, and the cells were collected and lysed by addition of 200 μl of radioimmunoprecipitation assay buffer containing protease and phosphatase inhibitors. Protein concentration of cell lysates was determined using BCA (Thermo Fisher Scientific). Samples were resolved under denaturing and reducing conditions using 4 to 12% bis-tris gels (NuPAGE) and transferred to a polyvinylidene fluoride membrane (Immobilon-P, Millipore). Membranes were blocked with 5% nonfat dried milk and incubated with anti-puromycin primary antibodies (1:1000 dilution, clone 12D10, Milipore Sigma) overnight at 4°C. After washing the membranes, secondary anti-mouse antibodies coupled with horseradish peroxidase were applied (Amersham-GE). Immunoreactive bands were visualized by enhanced chemiluminescence reagent (Thermo Fisher Scientific), and signal was acquired using ChemiDoc Imaging System (Bio-Rad).

**Monitoring of Protein Synthesis in Neurons**

Primary neurons were exposed or not to 10 ug/ml of cycloheximide for 30 minutes. Puroswitch was then added to the final concentration of 5 μM, and cells were incubated inside dark containers. To isolate the effects of light in the incorporation of puromycin, two containers were used: one containing DISCO light source, and one without. Cells were incubated in these conditions for 1h, with 390 nm light shined for 10 ms every 10 seconds. Media was then aspirated, cells were washed 1x with cold PBS and fixed in PBS + 4% paraformaldehyde for 20 min/RT, in the dark. Cells were washed 3x with PBS, and permeabilized with PBS + 0.5% Triton X-100 for 15 min/RT. After a quick rinse with PBS, coverslips were blocked with PBS + 5% normal goat serum, and then incubated overnight with two primary antibodies: mouse anti-puromycin (Millipore, clone 12D10, 1:500); and chicken anti-MAP2 (Encor Biotech, 1:1000). The next day, coverslips were washed 3x 5 min with PBS and incubated with two secondaries: goat anti-mouse 568 (Thermofisher, 1:500); and goat anti-chicken 647 (Thermofisher, 1:500). Coverslips were incubated in the dark for 1h and then washed 3x 5 min with PBS. The coverslips were then quickly rinsed in mili-q water and mounted in glass slides using Prolong antifade mounting media with DAPI (Thermofisher). For visualization, cells were imaged using a Leica SP8 upright microscope, using the 63X magnification objective. For each cell, a Z-stack containing 15 sections was obtained, with each stack separated 0.23 um from the next one. Maximum projection was obtained using Fiji/ImageJ. To generate a mask containing only the neuronal region, MAP2 staining was used. The region of interest (ROI) was then defined using this mask and the amount of diazo-puromycin overlapping with the ROI was measured. Raw Integrated Density (the sum of the intensity of all pixels within ROI) was normalized by the total area measured.

#### Synthetic Procedures and Characterization

Fmoc-phenylazophenylalanine (**1**)

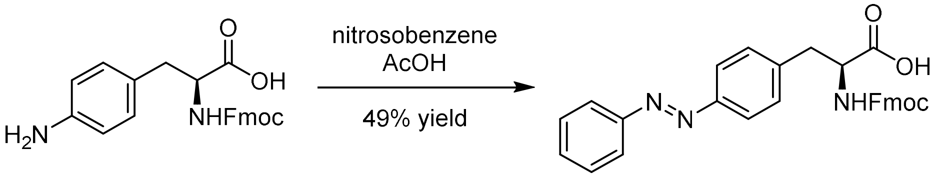


4-amino-N-Fmoc-L-phenylalanine (1.653 g, 4.108 mmol, 1.1 eq.), was dissolved in 30.8 ml of acetic acid. The solution was stirred until the substrate was fully dissolved and nitrosobenzene (0.4 g, 3.734 mmol, 1.0 eq.) was added to the reaction mixture. The solution was stirred overnight at room temperature and concentrated under reduced pressure. The resulting residue was dissolved in 100 ml of ethyl acetate and 100 ml of $H_2O$ was sequentially added. The aqueous layer was extracted (3 x 100 ml) with ethyl acetate. The organic layers were combined, washed with saturated aqueous sodium chloride (2 x 100 mL) and dried over sodium sulfate. Then the solvent was removed under reduced pressure and the residue purified by column chromatography (0 to 10% 10:1 methanol:acetic acid in dichloromethane) to afford the title compound (0.894 g, 1.819 mmol, 49%) as an orange solid.

**$R_f$** = 0.63 (10% methanol in dichloromethane)

**$^1$H-NMR** (400 MHz, DMSO-$d_6$) δ 7.88 – 7.85 (m, 4H), 7.76 (d, J = 8.0 Hz, 2H), 7.65 – 7.56 (m, 5H), 7.43 – 7.36 (m, 4H), 7.33 – 7.27 (m, 2H), 7.16 – 7.03 (m, 1H), 4.32 – 4.23 (m, 1H), 4.17 (q, J = 5.2, 3.4 Hz, 2H), 4.12 – 4.04 (m, 1H), 3.21 (dd, J = 13.6, 4.5 Hz, 1H), 3.00 (dd, J = 13.5, 8.7 Hz, 1H).

**$^{13}$C-NMR** (100 MHz, DMSO) δ 155.98, 152.46, 150.88, 144.35, 144.25, 141.14, 131.73, 130.82, 129.91, 128.02, 127.49, 125.70, 125.59, 122.86, 122.70, 120.54, 65.81, 56.64, 47.13, 40.62, 40.41, 40.20, 39.99, 39.78, 39.57, 39.36, 37.49.

| | | |
|---|---|---|
| **LCMS** (ESI): | $t_{ret}$ = 4.689 min | 492.1 m/z $[M+H]^+$. |
| **HRMS** (ESI): | calc. for $C_{30}H_{25}N_3O_4Na^+$: | 514.1737 m/z $[M+Na]^+$. |
| | found: | 514.1729 m/z $[M+Na]^+$. |

#### Azo-Puromycin

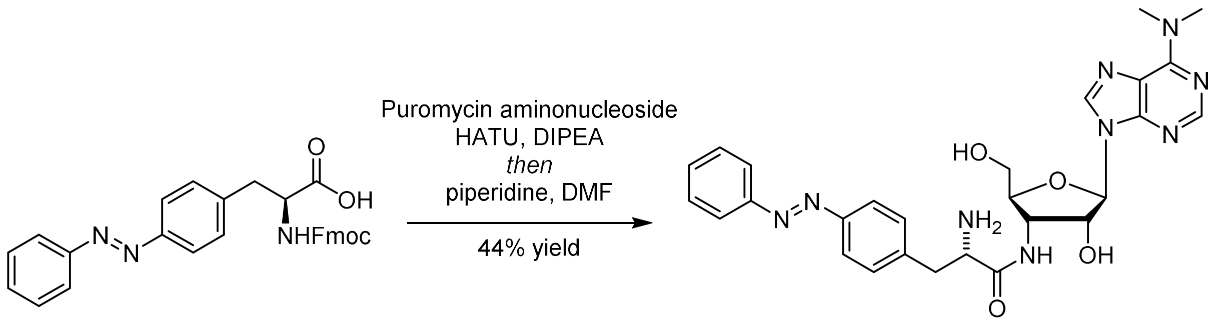


To puromycin aminonuleoside (10 mg, 0.034 mmol, 1.0 eq.), Fmoc-phenylazophenylalanine (**1**, 21.7 mg, 0.044 mmol, 1.3 eq.), HATU (19.4 mg, 0.051 mmol, 1.5 eq.) under nitrogen atmosphere was added anhydrous DMF (2.4 ml). The mixture was stirred until all solids were dissolved and distilled DIPEA (24 µl, 0.136 mmol, 4.0 eq) was added dropwise. The solution was stirred overnight at room temperature and concentrated under reduced pressure until most of the DMF had evaporated. The resulting residue was dissolved in DMF (0.107 ml) and piperidine (0.107 ml, 1.077 mmol, 31.7 eq.) was added dropwise. The solution was stirred for one hour at room temperature and concentrated under reduced pressure. The resulting residue was dissolved in 1 ml of ethyl acetate and 1 ml of 10% LiCl aqueous solution. Subsequently, the aqueous layer was extracted (3 x 1 ml) with ethyl acetate. The organic layers were combined, washed with saturated aqueous sodium chloride (1 mL) and dried over sodium sulfate. The solvent was removed under reduced pressure and the residue purified by column chromatography (0 to 10% methanol in dichloromethane) to afford the title compound (8.3 mg, 0.015 mmol, 44%) as a light orange solid.

**$R_f$** = 0.57 (10% methanol in dichloromethane)

**$^1$H NMR** (600 MHz, MeOD-*d*$_4$) δ 8.30 (s, 1H), 8.17 (s, 1H), 7.88 – 7.83 (m, 4H), 7.54 – 7.46 (m, 3H), 7.42 (d, J = 8.3 Hz, 2H), 5.92 – 5.89 (m, 1H), 4.58 – 4.51 (m, 3H), 4.01 – 3.92 (m, 1H), 3.81 – 3.76 (m, 1H), 3.72 (t, J = 7.1 Hz, 1H), 3.54 (dd, J = 12.5, 3.1 Hz, 1H), 3.47 (s, 8H), 3.13 – 3.03 (m, 1H), 3.00 – 2.89 (m, 1H).

**$^{13}$C-NMR** (100 MHz, MeOD) δ 176.69, 156.15, 153.98, 152.99, 152.96, 152.91, 150.57, 142.55, 139.16, 132.21, 131.36, 130.24, 124.01, 123.74, 91.94, 91.90, 84.89, 84.85, 75.11, 75.08, 62.23, 54.80, 51.93, 49.64, 49.43, 49.21, 49.00, 48.79, 48.57, 48.36, 42.26, 39.03.

| | | |
|---|---|---|
| **LCMS** (ESI): | $t_{ret}$ = 2.61 min | 546.2 m/z $[M+H]^+$. |
| **HRMS** (ESI): | calc. for $C_{27}H_{32}N_9O_4^+$: | 546.2572 m/z $[M+H]^+$. |
| | found: | 546.2436 m/z $[M+H]^+$. |

Phthalimide (**S2**)

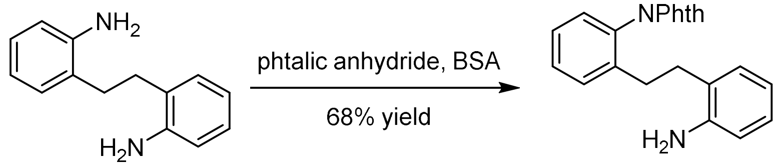


To a warm solution of 2,2'-ethylenedianiline (**S1**, 10.00 g, 47.1 mmol, 1.0 eq.) in toluene (94.2 mL) was added bis(trimethylsilyl)acetamide (11.6 mL, 47.1 mmol, 1.0 eq.), followed by phthalic anhydride (6.98 g, 47.1 mmol, 1.0 eq.). The mixture was stirred overnight at 100 °C, then cooled to room temperature and concentrated under reduced pressure. Dichloromethane (120 mL) was added to the residue and the mixture was washed with 2 M aqueous sodium hydroxide (100 mL) and water (2×100 mL). After drying over sodium sulfate, the resulting solution was concentrated under reduced pressure. Diethyl ether (100 mL) was added to the oily residue and the mixture was sonicated until the oily layer had fully dissolved and large amounts of a yellow solids had formed, then the mixture was cooled with an ice-bath for two hours. Next, the solids were collected and washed with a small amount of additional diethyl ether. Drying of the solids under reduced pressure afforded the title compound (10.92 g, 31.89 mmol, 68%) as a yellow solid.

$\mathbf{R_f}$ = 0.88 (20% ethyl acetate in dichloromethane).

**$^1$H-NMR** (400 MHz, $CDCl_3$): 7.95 (dd, J = 5.3, 3.1 Hz, 2H), 7.81 (dd, J = 5.4, 3.1 Hz, 2H), 7.39 (m, J = 21.7, 7.5 Hz, 3H), 7.20 (d, J = 7.1 Hz, 1H), 7.02 – 6.95 (m, 2H), 6.66 (t, J = 7.4 Hz, 1H), 6.56 (d, J = 8.1 Hz, 1H), 2.84 – 2.71 (m, 4H).

**$^{13}$C-NMR** (100 MHz, $CDCl_3$): 168.10, 144.43, 140.34, 134.56, 132.13, 130.52, 130.22, 129.83, 129.75, 129.35, 127.46, 127.38, 125.50, 123.96, 118.85, 115.76, 32.25, 31.23, 31.09.

| | | |
|---|---|---|
| **LCMS** (ESI): | $t_{ret}$ = 3.42 min | 343.2 m/z $[M+H]^+$. |
| **HRMS** (ESI): | calc. for $C_{22}H_{19}N_2O_2^+$: | 343.1441 m/z $[M+H]^+$. |
| | found: | 343.1433 m/z $[M+H]^+$. |

2-(2-aminophenethyl)-4-bromoaniline (**S3**)

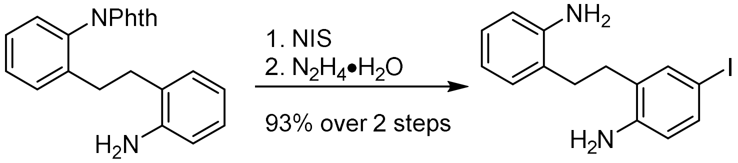


**Iodination:** To a solution of mono-phthalimide (**S2**, 2.50 g, 7.30 mmol, 1.0 eq.) in anhydrous dimethyl sulfoxide (21 mL) was added N-iodosuccinimide (1.64 g, 7.30 mmol, 1.0 eq.) in one portion. After the mixture had stirred for three hours, saturated, aqueous sodium chloride/water = 1/4 (50 mL) was added and the mixture was extracted with dichloromethane (30 mL and 2 x 15 mL). The extracts were washed with saturated, aqueous sodium chloride/water = 1/4 (2×50 mL) and dried over sodium sulfate. The solution was then filtered over a short plug of silica (5 g) and eluted with additional dichloromethane (25 mL). The solvent was removed under reduced pressure and the residue was used without further purification.

**Phthalimide cleavage:** To the iodination product were added tetrahydrofuran (37 mL) and hydrazine monohydrate (2.13 mL, 43.8 mmol, 6 eq.) and the mixture was stirred at reflux for two hours. After cooling to room temperature, the mixture was filtered. The solids were washed with additional tetrahydrofuran (2 x 10 mL) and discarded. Water (20 mL) was added to the filtrate, the tetrahydrofuran removed under reduced pressure and the residue extracted with diethyl ether (30 mL and 2 x 15 mL). The extracts were washed with saturated, aqueous sodium chloride (30 mL), dried over sodium sulfate and the solvent was removed under reduced pressure. The residue was purified by precipitation from dichloromethane solution by addition of hexanes to afford the title compound (1.98 g, 6.80 mmol, 93%) as a brownish-yellow solid.

$\mathbf{R_f}$ = 0.38 (20% ethyl acetate in dichloromethane).

**$^{1}$H-NMR** (400 MHz, $CDCl_3$): 7.36 (s, 1H), 7.31 (d, J = 8.3 Hz, 1H), 7.10 – 7.01 (m, 2H), 6.76 (t, J = 7.4 Hz, 1H), 6.70 (d, J = 7.8 Hz, 1H), 6.45 (d, J = 8.3 Hz, 1H), 3.44 (s, 4H), 2.82 – 2.68 (m, 4H).

**$^{13}$C-NMR** (100 MHz, $CDCl_3$): 144.29, 144.22, 137.99, 135.99, 129.71, 129.09, 127.62, 126.01, 119.39, 117.99, 116.18, 80.22, 31.15, 30.70.

| | | |
|---|---|---|
| **LCMS** (ESI): | $t_{ret}$ = 3.41 min | 339.0 m/z $[M+H]^+$. |
| **HRMS** (ESI): | calc. for $C_{14}H_{16}IN_2^+$: | 339.0353 m/z $[M+H]^+$. |
| | found: | 339.0351 m/z $[M+H]^+$. |

(Z)-2-iodo-11,12-dihydrodibenzo[c,g][1,2]diazocine (**3**)

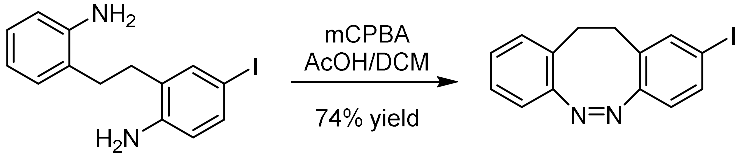


A freshly prepared and titrated solution of mCPBA (1.02 g, 5.91 mmol, 2.0 eq.) in acetic acid (10 ml) was added by syringe pump within a period of 12 hours under rapid stirring to a solution of a 2-(2-aminophenethyl)-4-iodoaniline (**S3**, 1.00 g, 2.96 mmol, 1.0 eq.) in acetic acid/dichloromethane = 1/3 (75.7 mL). After the complete addition of the mCPBA solution, the mixture was stirred for at least one more hour. The solvent was then removed under reduced pressure, the residue taken up in ethyl acetate (70 mL) and washed with saturated, aqueous sodium hydrogen carbonate (2 x 70 mL), followed by saturated, aqueous sodium chloride (70 mL). The solution was dried over sodium sulfate, the solvent was removed under reduced pressure. The crude product was then purified by column chromatography (0 to 10% ethyl acetate in hexanes) to afford the title compound (0.73 g, 2.19 mmol, 74%) as a yellow solid.

**$R_f$** = 0.43 (10% ethyl acetate in hexanes).

**$^1$H-NMR** (400 MHz, $CDCl_3$): 7.44 (dd, J = 8.3, 1.8 Hz, 1H), 7.34 (d, J = 1.8 Hz, 1H), 7.16 (td, J = 7.5, 1.5 Hz, 1H), 7.05 (td, J = 7.5, 1.3 Hz, 1H), 7.00 (dd, J = 7.6, 1.5 Hz, 1H), 6.83 (dd, J = 7.8, 1.3 Hz, 1H), 6.58 (d, J = 8.2 Hz, 1H), 3.07 – 2.61 (m, 4H).

**$^{13}$C-NMR** (100 MHz, $CDCl_3$): 155.4, 155.0, 138.4, 135.8, 130.7, 129.9, 127.6, 127.5, 127.1, 120.9, 118.9, 91.9, 31.6, 31.5.

| | | |
|---|---|---|
| **LCMS** (ESI): | $t_{ret}$ = 4.36 min | 335.0 m/z $[M+H]^+$. |
| **HRMS** (ESI): | calc. for $C_{14}H_{12}IN_2^+$: | 335.0040 m/z $[M+H]^+$. |
| | found: | 335.0051 m/z $[M+H]^+$. |

methyl (*S*,*Z*)-2-((*tert*-butoxycarbonyl)amino)-3-(11,12-dihydrodibenzo[*c*,*g*][1,2]diazocin-2-yl)propanoate (**5**)

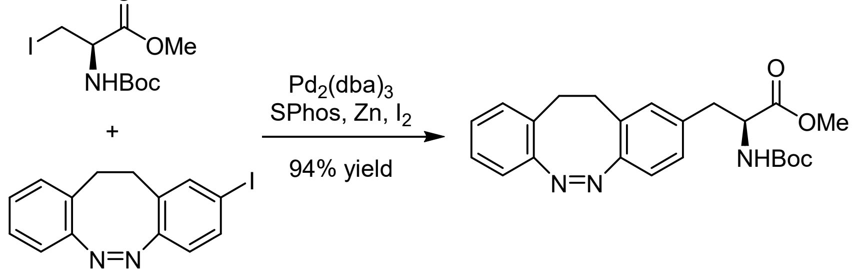


**Preparation of zinc dust:** To a 250 mL round bottom flask was added, 100 mL of 2% HCl acid and zinc powder (16.0 g, 0.245 mol). The solution was stirred vigorously, until the surface of the zinc becomes bright. The aqueous solution was then decanted, and the zinc powder in the flask was washed by decantation with four 200 mL portions of water. The zinc powder was filtered with 200 mL of water and washed successively with 50 mL of ethanol, 100 mL of acetone, and 50 mL of diethyl ether. Filtration and washing were done as quickly as possible to avoid exposure to air. The resulting zinc was transferred to an oven dried round-bottom flask and dried overnight under reduced atmosphere pressure. The flask was refilled with nitrogen and stored at room temperature until use.

**Negishi Cross coupling:** To the prepared zinc dust (0.6 g, 9.12 mmol, 3 eq.) and iodide (0.12 g, 0.46 mmol, 0.15 eq.) under nitrogen atmosphere was added anhydrous DMF (3 ml). The suspension was stirred until a color change from purple to clear was observed. N-Boc-L-iodoalaninemethylester (1.00 g, 3.04 mmol, 1.0 eq.) and additional iodide (0.12 g, 0.46 mmol, 0.15 eq.) were sequentially added. The mixture was stirred for 30 minutes where a color change from purple to clear and an exotherm were noticed. The solution was filtered through a teflon syringe filter into another flask under nitrogen athmosphere. $Pd_2dba_3$ (70 mg, 276µmol, 0.025 eq.), SPhos (62 mg, 0.15 mmol, 0.05 eq.) and (Z)-2-iodo-11,12-dihydrodibenzo[c,g][1,2]diazocine (**3**, 1.32 g, 3.95 mmol, 1.3 eq.) were added to the filtered solution and stirred overnight. The stir bar was removed and Celite® added until the mixture has the consistency of damp sand. The reaction flask was dried under reduced pressure and the resulting powder directly purified by column chromatography (0 to 80% ethyl acetate in hexanes) to afford the title compound (1.17 g, 2.86 mmol, 94%) as a yellow oil.

$\mathbf{R_f}$ = 0.51 (40% ethyl acetate in hexanes)

**$^1$H-NMR** (400 MHz, $CDCl_3$): 7.13 (t, J = 7.4 Hz, 1H), 7.02 (d, J = 7.5 Hz, 1H), 6.98 (t, J = 6.5 Hz, 1H), 6.89 (m, 1H), 6.82 (d, J = 7.8 Hz, 1H), 6.74 (m, J = 18.7 Hz, 2H), 4.88 (s, 1H), 4.46 (d, J = 6.7 Hz, 1H), 3.53 (s, 3H), 2.93 (m, J = 19.4 Hz, 4H), 2.83 – 2.63 (m, 2H), 1.41 (s, 9H).

**$^{13}$C-NMR** (100 MHz, $CDCl_3$): 172.28, 155.63, 155.08, 154.52, 135.07, 130.66, 129.68, 128.42, 128.15, 127.62, 127.16, 126.84, 119.22, 118.89, 80.15, 54.47, 52.22, 38.11, 31.98, 31.69, 28.44.

| | | |
|---|---|---|
| **LCMS** (ESI): | $t_{ret}$ = 4.18 min | 410.2 m/z $[M+H]^+$. |
| **HRMS** (APCI): | calc. for $C_{23}H_{28}N_3O_4^+$: | 410.2074 m/z $[M+H]^+$. |
| | found: | 410.2078 m/z $[M+H]^+$. |

(S,Z)-2-((tert-butoxycarbonyl)amino)-3-(11,12-dihydrodibenzo[c,g][1,2]diazocin-2-yl)propanoic acid (**S4**)

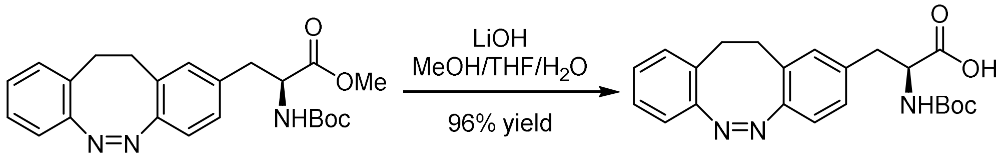


To a solution of methyl (*S*,*Z*)-2-((*tert*-butoxycarbonyl)amino)-3-(11,12-dihydrodibenzo[*c*,*g*][1,2]diazocin-2-yl)propanoate (**5**, 0.519 g, 1.27 mmol, 1.0 eq.) in tetrahydrofuran (3.6 ml), methanol (3.6 ml) and water (3.6 ml) was added lithium hydroxide (0.085 g, 3.55 mmol, 2.8 eq.) and stirred overnight. The mixture was concentrated under reduced pressure until about 20% of the solution remained. The solution was acidified with 2M HCl until the pH was around 1. To the resulting suspension 10 ml of dichloromethane was added and the precipitate dissolved. The organic layer was then extracted and the aqueous was washed (3 x 5 ml) with dichloromethane until the aqueous layer was clear. The organic layers were combined, washed with saturated aqueous sodium chloride (10 mL) and dried over sodium sulfate. The solvent was removed under reduced pressure to afford the title compound as a yellow solid (0.478 g, 1.21 mmol, 96%), which was used without purification in the next step.

**$R_f$** = 0.47 (5% methanol in dichloromethane)

**$^1$H-NMR** (400 MHz, DMSO-$d_6$) δ 12.59 (s, 1H), 7.16 (t, *J* = 7.4, 1.7 Hz, 1H), 7.10 – 7.01 (m, 3H), 6.93 (s, 1H), 6.83 (d, *J* = 7.8, 1.2 Hz, 1H), 6.76 (d, *J* = 8.0 Hz, 1H), 4.02 (td, *J* = 9.4, 9.0, 4.4 Hz, 1H), 2.93 – 2.60 (m, 6H), 1.31 (s, 9H).

**$^{13}$C-NMR** (100 MHz, DMSO) δ 173.39, 155.30, 155.08, 153.45, 136.94, 130.47, 130.13, 129.69, 127.81, 127.22, 127.06, 126.75, 118.61, 118.37, 77.99, 54.71, 35.73, 31.29, 30.72, 28.21, 28.15, 27.82.

**LCMS** (ESI): $t_{ret}$ = 3.70 min 396.2 m/z $[M+H]^+$.

**HRMS** (ESI): calc. for $C_{22}H_{25}N_3NaO_4^+$: 418.1737 m/z $[M+Na]^+$.
found: 418.174 m/z $[M+Na]^+$.

(S,Z)-2-((((9H-fluoren-9-yl)methoxy)carbonyl)amino)-3-(11,12-dihydrodibenzo[c,g][1,2]diazocin-2-yl)propanoic acid (**6**)

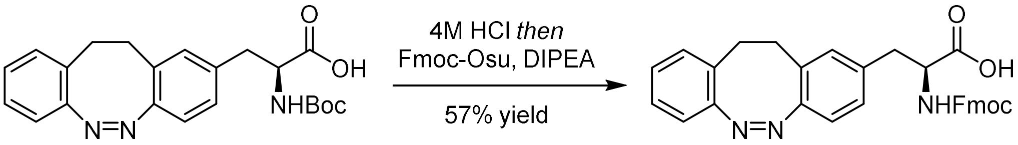


To a solution of (S,Z)-2-((tert-butoxycarbonyl)amino)-3-(11,12-dihydrodibenzo [c,g][1,2]diazocin-2-yl)propanoic acid (**S4**, 0.100 g, 0.253 mmol, 1.0 eq.) in dichloromethane (5.2 ml) was added 4M HCl in dioxane (1.6 ml) dropwise and stirred overnight. The yellow solids that precipitated were sonicated into the solution and the mixture was concentrated under reduced pressure. The solids were sequentially suspended in dichloromethane (2.5 ml) and Fmoc-Osu (0.128 g, 0.379 mmol, 1.5 eq.) was added to the mixture. Under ice-bath cooling, N,N-Diisopropylethylamine (0.088 ml, 0.506 mmol, 2.0 eq.) was added dropwise and the pH was checked to ensure that the solution was basic. The solution was then stirred for 2 hours and the solution washed with 2M HCl (3 x 5 ml), saturated aqueous sodium chloride (5 mL) and dried over sodium sulfate. The solvent was removed under reduced pressure and the residue purified by column chromatography (0 to 10% methanol in dichloromethane) to afford the title compound (0.075 g, 0.144 mmol, 57%) as a yellow solid.

**$R_f$** = 0.42 (5% methanol in dichloromethane)

**$^1$H-NMR** (400 MHz, DMSO-$d_6$) $\delta$ 12.68 (s, 1H), 7.90 (d, *J* = 7.6 Hz, 2H), 7.74 – 7.59 (m, 3H), 7.42 (t, *J* = 7.5 Hz, 2H), 7.32 (d, *J* = 7.5 Hz, 2H), 7.18 – 6.89 (m, 4H), 6.85 – 6.71 (m, 2H), 4.31 – 4.05 (m, 4H), 2.99 – 2.86 (m, 1H), 2.84 – 2.53 (m, 5H).

**$^{13}$C-NMR** (100 MHz, DMSO) δ 173.15, 155.83, 155.06, 153.48, 143.72, 140.66, 136.85, 130.18, 129.66, 127.77, 127.63, 127.05, 126.70, 125.26, 120.10, 118.39, 65.60, 55.16, 46.55, 35.78, 31.23, 30.70.

**LCMS** (ESI): $t_{ret}$ = 4.24 min 518.2 m/z $[M+H]^+$.

**HRMS** (ESI): calc. for $C_{32}H_{27}N_3NaO_4^+$: 540.1894 m/z $[M+Na]^+$.
found: 540.1882 m/z $[M+Na]^+$.

Fmoc-puroswitch (**7**)

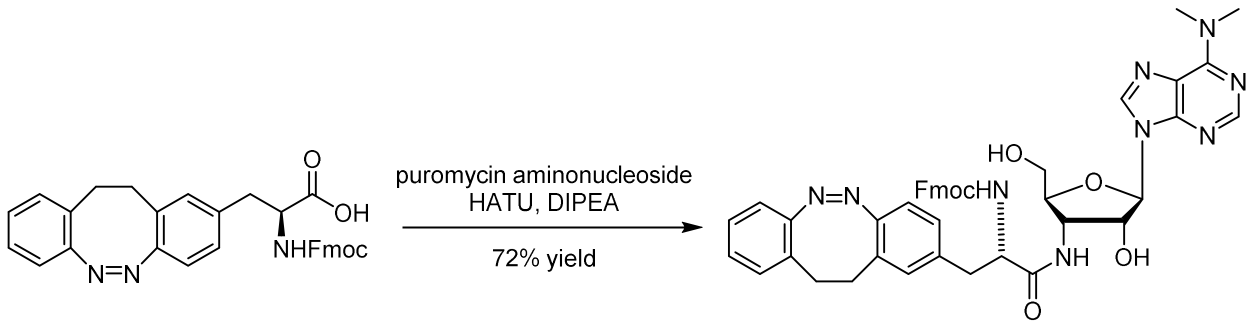


To puromycin aminonuleoside (20 mg, 0.068 mmol, 1.0 eq.), (S,Z)-2-((((9H-fluoren-9-yl)methoxy)carbonyl)amino)-3-(11,12-dihydrodibenzo[c,g][1,2]diazocin-2-yl)propanoic acid (**6**, 52.8 mg, 0.102 mmol, 1.5 eq.), HATU (38.8 mg, 0.102 mmol, 1.5 eq.) under nitrogen atmosphere was added anhydrous DMF (4.9 ml). The mixture was stirred until all solids were dissolved and distilled DIPEA (47 μl, 0.272 mmol, 4.0 eq) was added dropwise. The solution was stirred overnight at room temperature and concentrated under reduced pressure until most of the DMF had evaporated. The resulting residue was dissolved in 5 ml of dichloromethane and 5 ml of 10% LiCl aqueous solution were added. Subsequently, the layers were separated, and the aqueous layer was further extracted (3 x 5 ml) with dichloromethane. The organic layers were combined, washed with saturated aqueous sodium chloride (5 mL) and dried over sodium sulfate. The solvent was removed under reduced pressure and the residue purified by column chromatography (0 to 10% methanol in dichloromethane) to afford the title compound (39 mg, 0.049 mmol, 72%) as a yellow-brown solid.

$\mathbf{R_f}$ = 0.5 (1.25% methanol in dichloromethane)

**$^1$H-NMR** (400 MHz, DMSO-$d_6$) δ 8.44 (s, 1H), 8.23 (s, 1H), 8.16 (d, *J* = 7.3 Hz, 1H), 7.89 (d, *J* = 7.6 Hz, 2H), 7.74 – 7.48 (m, 3H), 7.42 (td, *J* = 7.5, 4.4 Hz, 2H), 7.35 – 7.28 (m, 2H), 7.16 – 7.1 (m, 1H), 7.08 – 6.94 (m, 2H), 6.91 (d, *J* = 4.4 Hz, 1H), 6.83 – 6.67 (m, 2H), 6.10 (s, 1H), 5.99 (d, *J* = 2.4 Hz, 1H), 4.47 (dd, *J* = 12.9, 5.6 Hz, 2H), 4.32 (dq, *J* = 10.5, 6.2, 5.0 Hz, 1H), 4.11 (d, *J* = 12.7 Hz, 3H), 3.93 (d, *J* = 4.9 Hz, 1H), 3.67 (d, *J* = 12.6 Hz, 1H), 3.52 – 3.38 (m, 7H), 2.89 – 2.61 (m, 6H).

**$^{13}$C-NMR** (100 MHz, DMSO) δ 171.69, 155.64, 155.07, 154.17, 153.42, 151.74, 149.61, 143.71, 140.66, 137.90, 136.90, 130.74, 130.32, 129.62, 127.92, 127.78, 127.62, 127.35, 127.20, 127.04, 126.94, 126.66, 125.33, 120.08, 119.60, 118.30, 89.37, 83.36, 73.03, 65.69, 60.83, 55.79, 50.27, 46.51, 37.32, 31.29, 30.72.

| | | |
|---|---|---|
| **LCMS** (ESI): | $t_{ret}$ = 3.95 min | 794.3 m/z $[M+H]^+$. |
| **HRMS** (ESI): | calc. for $C_{44}H_{44}N_9O_6^+$: | 794.3409 m/z $[M+H]^+$. |
| | found: | 794.3414 m/z $[M+H]^+$. |

#### Puroswitch

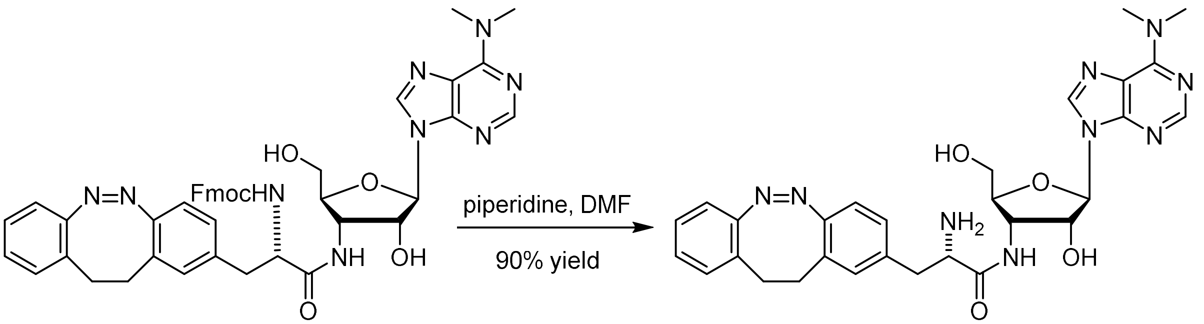

To a solution of Fmoc-puroswitch (**7**, 29.6 mg, 0.037 mmol, 1.0 eq.) in DMF (0.75 ml) was added piperidine (0.143 ml, 1.447 mmol, 38.8 eq.) dropwise. The solution was stirred for one hour at room temperature and concentrated under reduced pressure. The resulting residue was dissolved in 3 ml of ethyl acetate and 5 ml of 10% LiCl aqueous solution. Subsequently, the layers were separated, and the aqueous layer was further extracted (3 x 3 ml) with dichloromethane. The organic layers were combined, washed with saturated aqueous sodium chloride (5 mL) and dried over sodium sulfate. The solvent was removed under reduced pressure and the residue purified by column chromatography (0 to 7% methanol in dichloromethane) to afford the title compound (19.1 mg, 0.033 mmol, 90%) as a light-yellow solid.

**$R_f$** = 0.57 (10% methanol in dichloromethane)

**$^1$H-NMR** (600 MHz, DMSO-$d_6$) δ 8.44 (s, 1H), 8.23 (s, 1H), 8.05 (s, 1H), 7.16 (t, $J$ = 7.6, 1.5 Hz, 1H), 7.10 – 7.02 (m, 3H), 6.94 (d, $J$ = 8.1 Hz, 1H), 6.83 (d, $J$ = 7.8 Hz, 1H), 6.76 (t, $J$ = 6.8 Hz, 1H), 6.12 (s, 1H), 5.98 (d, $J$ = 2.8 Hz, 1H), 5.14 (s, 1H), 4.54 – 4.38 (m, 2H), 4.01 – 2.97 (s, 6H), 3.96 – 3.88 (m, 1H), 3.73 – 3.64 (m, 1H), 3.52 – 3.45 (m, 1H), 3.43 – 3.38 (m, 1H), 2.82 (d, $J$ = 7.5 Hz, 1H), 2.81 – 2.71 (m, 4H), 2.47 – 2.43 (m, 1H).

**$^{13}$C-NMR** (150 MHz, DMSO) δ 174.71, 155.14, 154.26, 153.36, 151.86, 149.63, 137.94, 137.76, 130.75, 130.36, 129.71, 127.93, 127.80, 127.46, 127.07, 126.75, 119.63, 118.41, 89.44, 83.52, 73.19, 60.94, 55.90, 50.02, 40.32, 40.06, 31.27, 30.72.

**LCMS** (ESI): $t_{ret}$ = 2.45 min 572.3 m/z $[M+H]^+$.

**HRMS** (ESI): calc. for $C_{29}H_{34}N_9O_4^+$: 572.2728 m/z $[M+H]^+$.
found: 572.2737 m/z $[M+H]^+$.

$^{1}$H-NMR (400 MHz, $CDCl_3$) – **1**

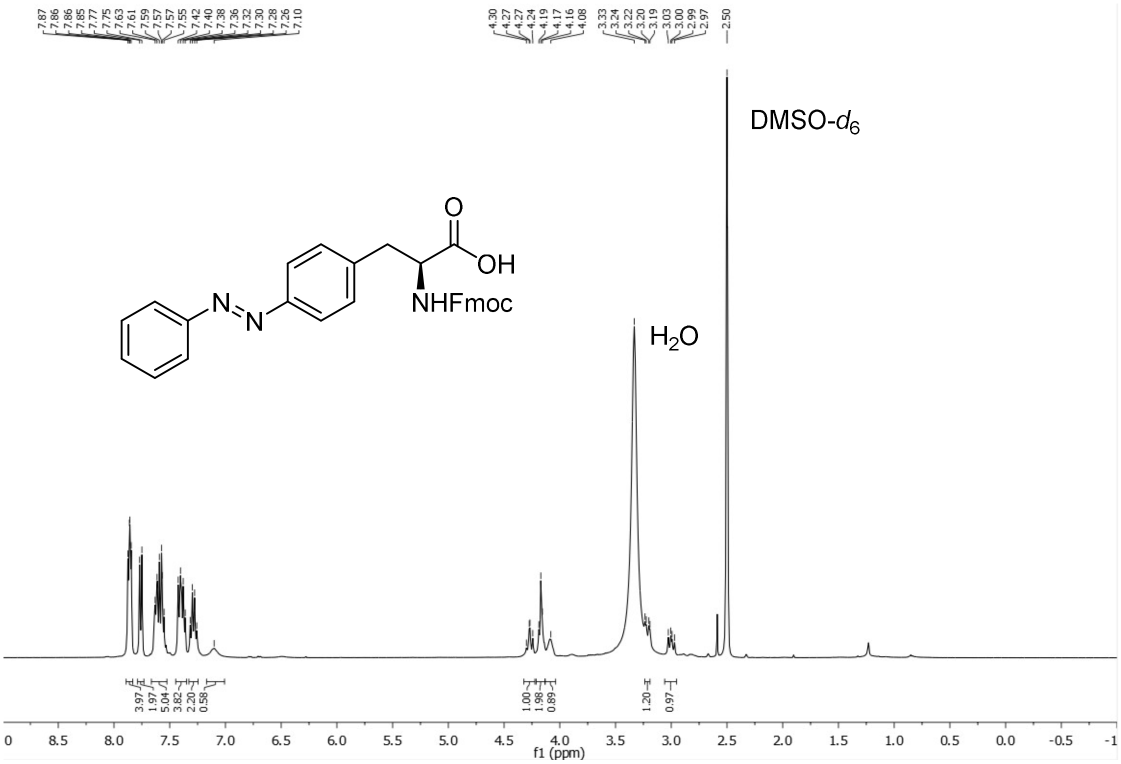


$^{13}$C-NMR (100 MHz, $CDCl_3$) – **1**

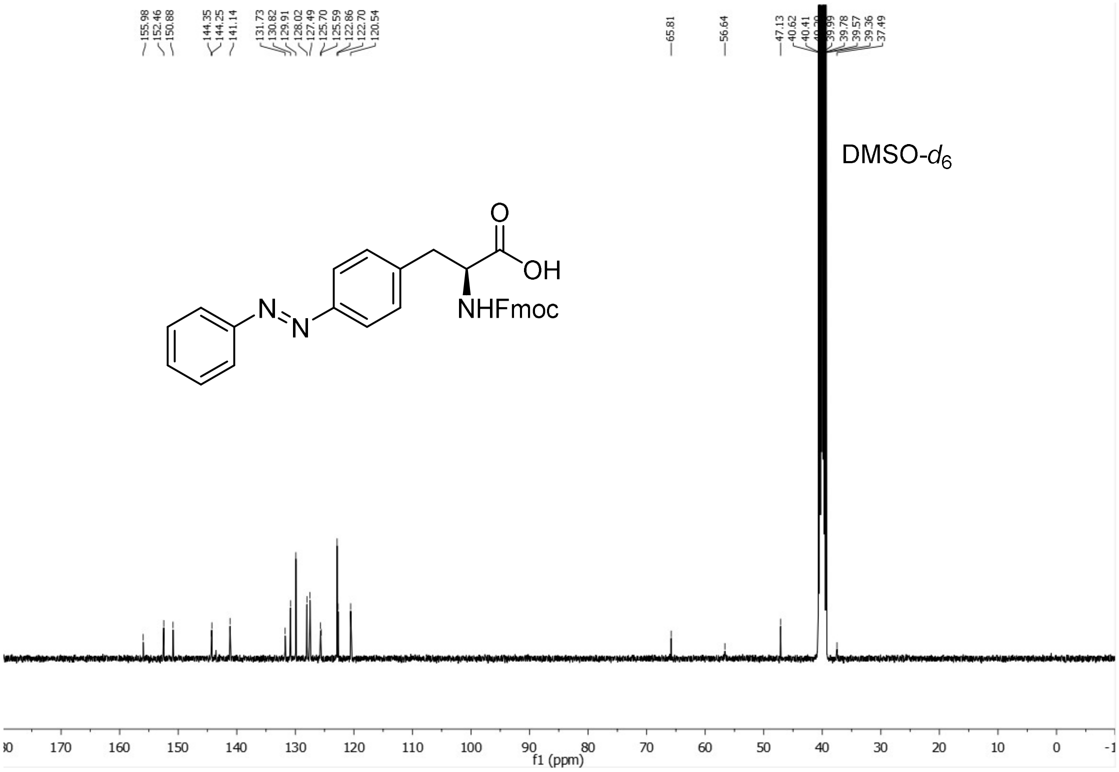

$^1$H-NMR (400 MHz, $CDCl_3$) – **azo-puromycin**

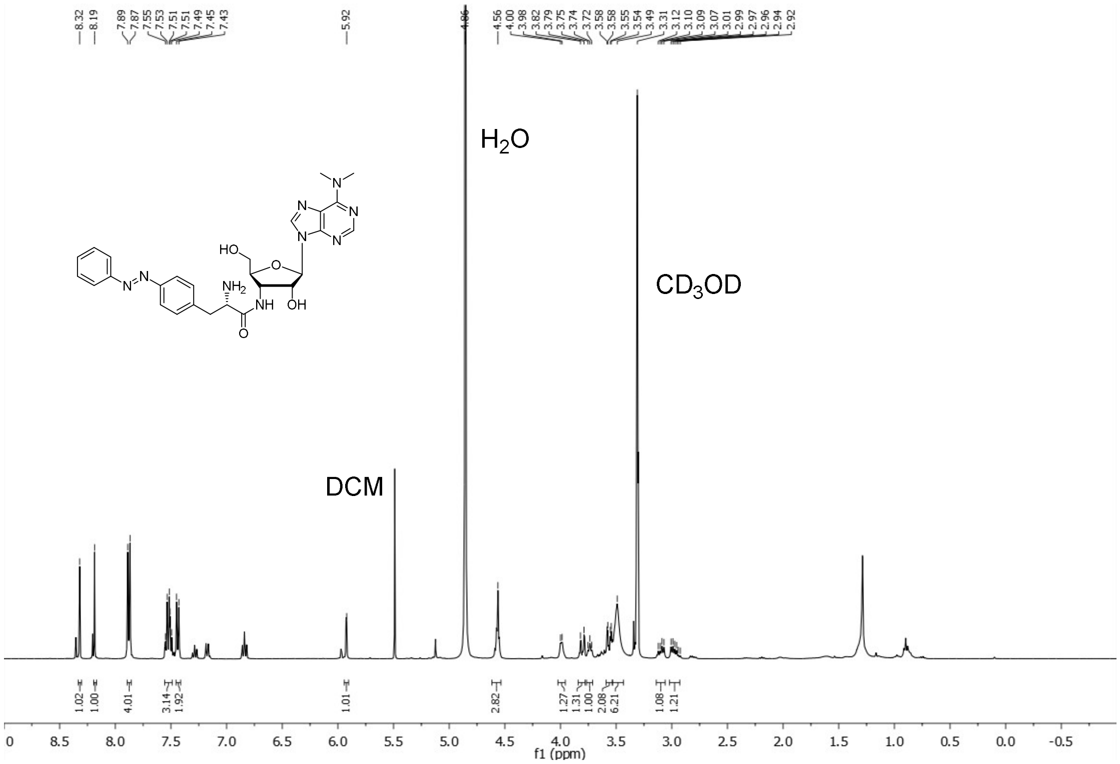


$^{13}$C-NMR (100 MHz, $CDCl_3$) – **azo-puromycin**

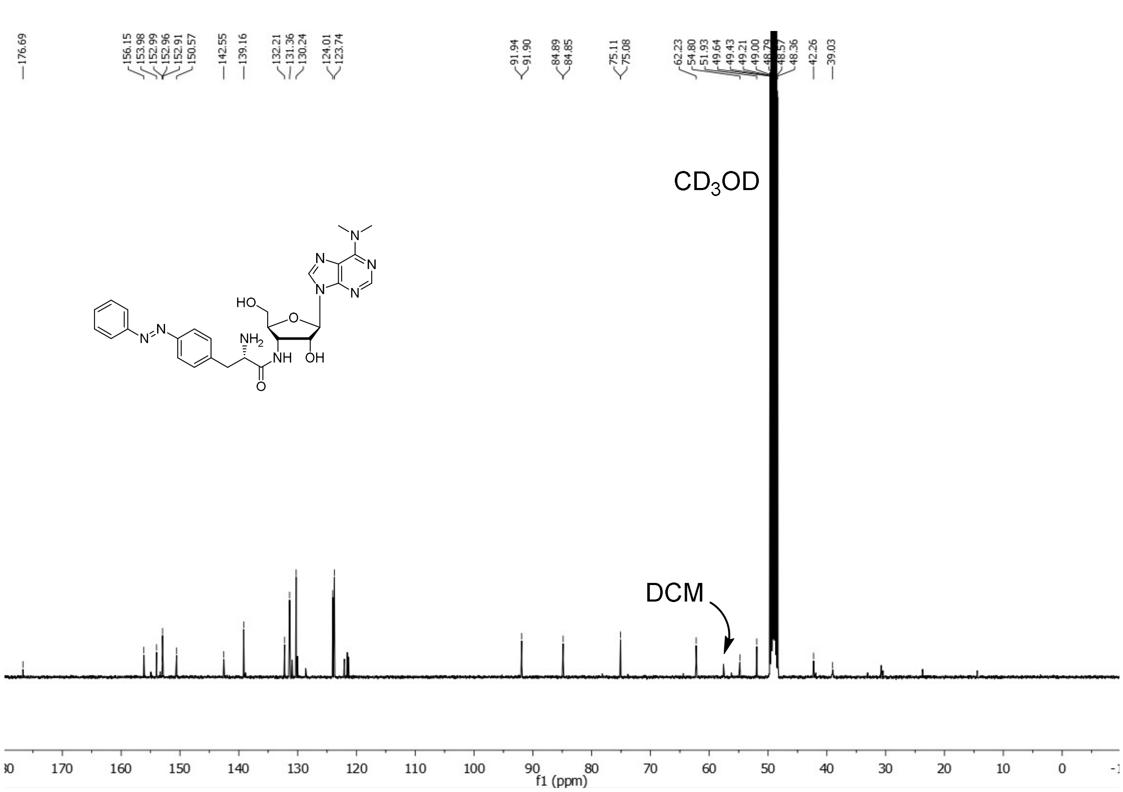

$^{1}$H-NMR (400 MHz, $CDCl_3$) – **S2**

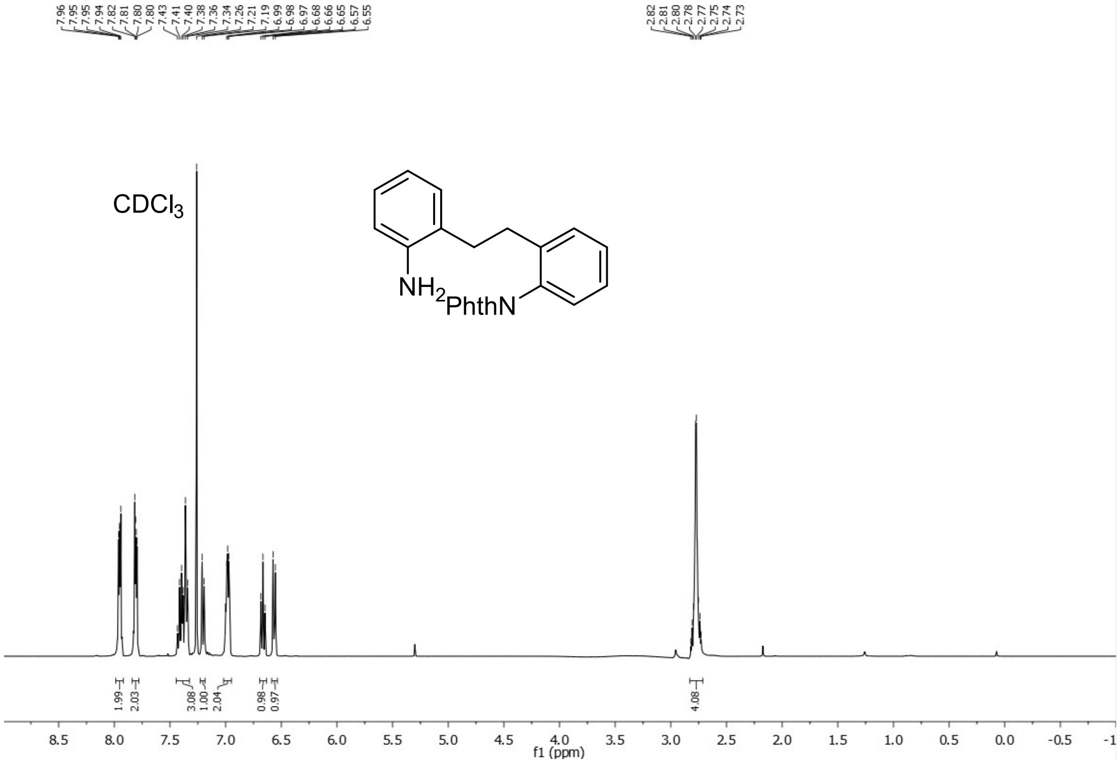


$^{13}$C-NMR (100 MHz, $CDCl_3$) – **S2**

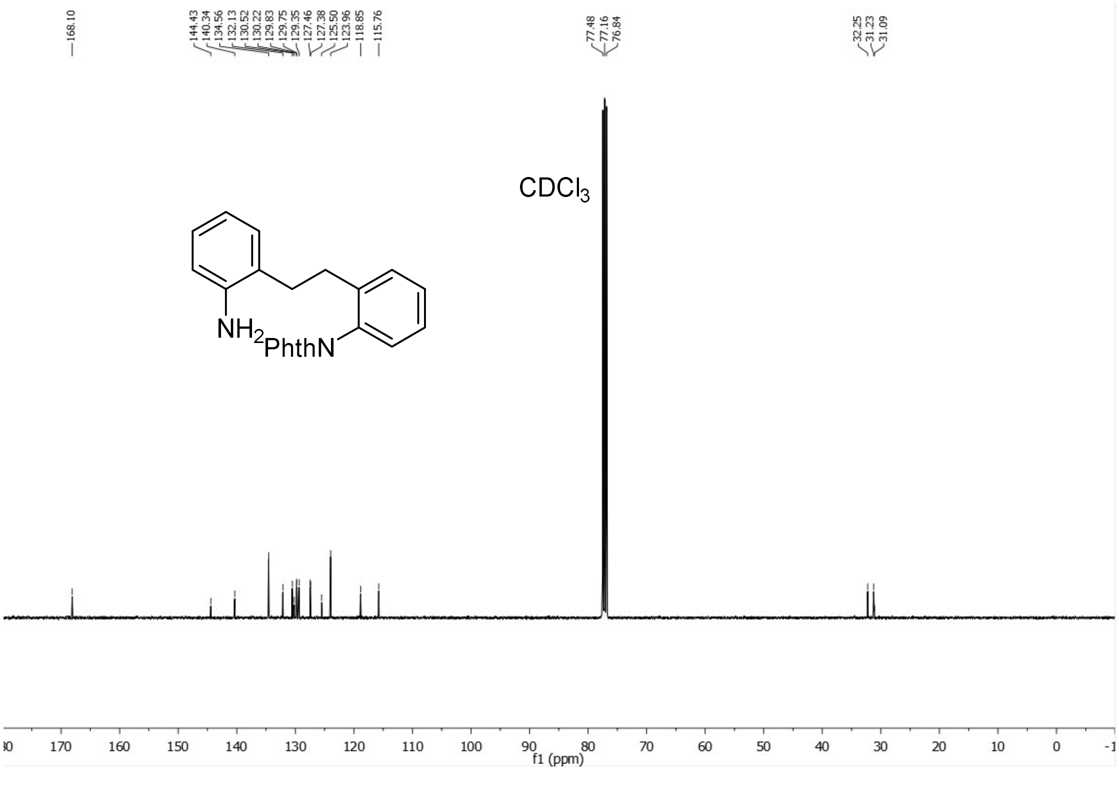

$^{1}$H-NMR (400 MHz, $CDCl_3$) – **S3**

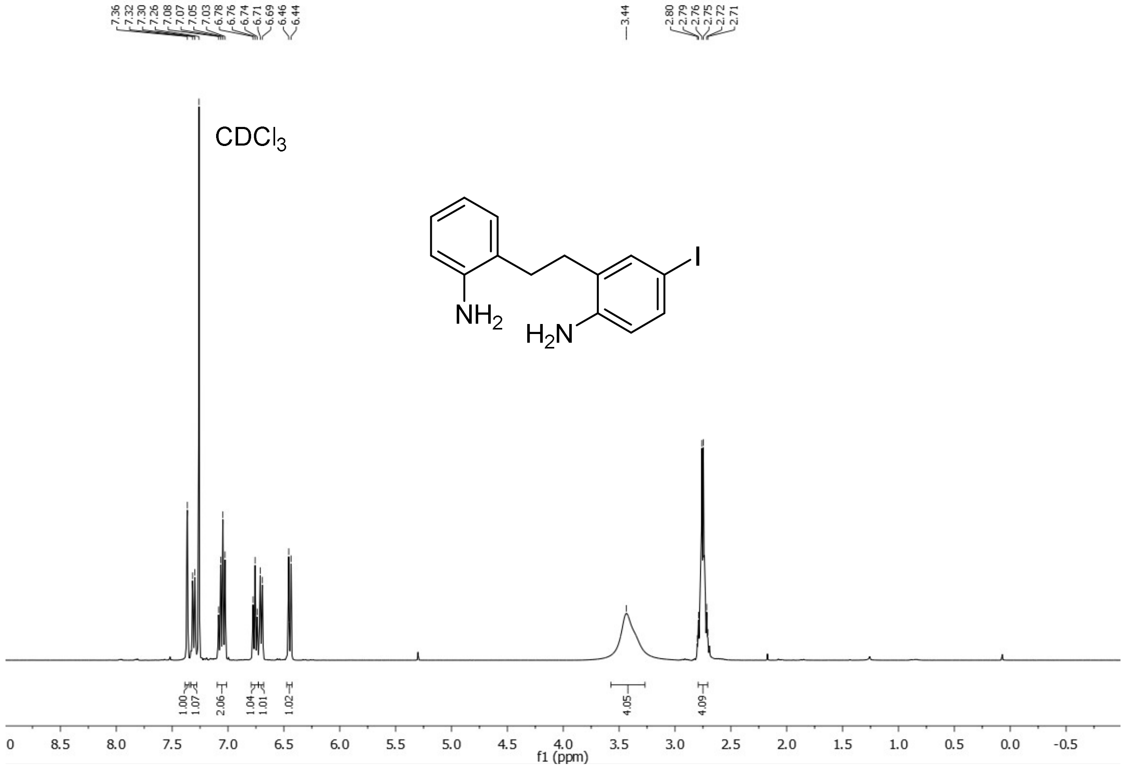


$^{13}$C-NMR (100 MHz, $CDCl_3$) – **S3**

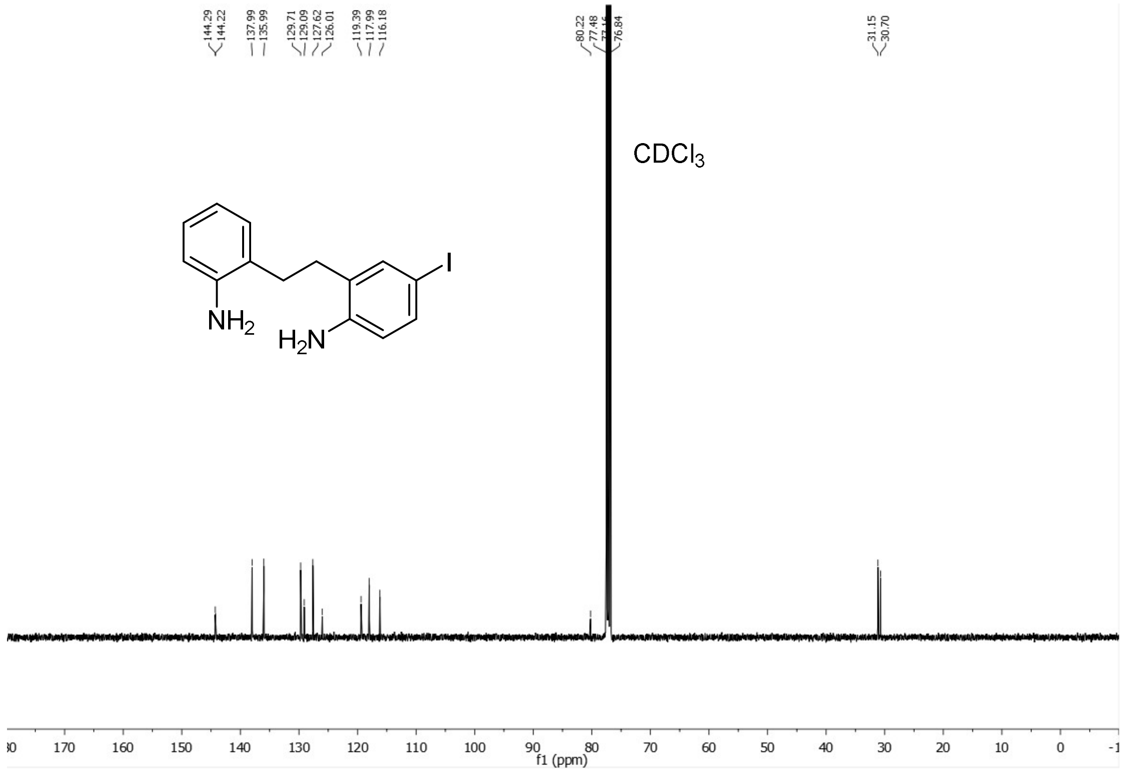

$^{1}$H-NMR (400 MHz, $CDCl_3$) – **3**

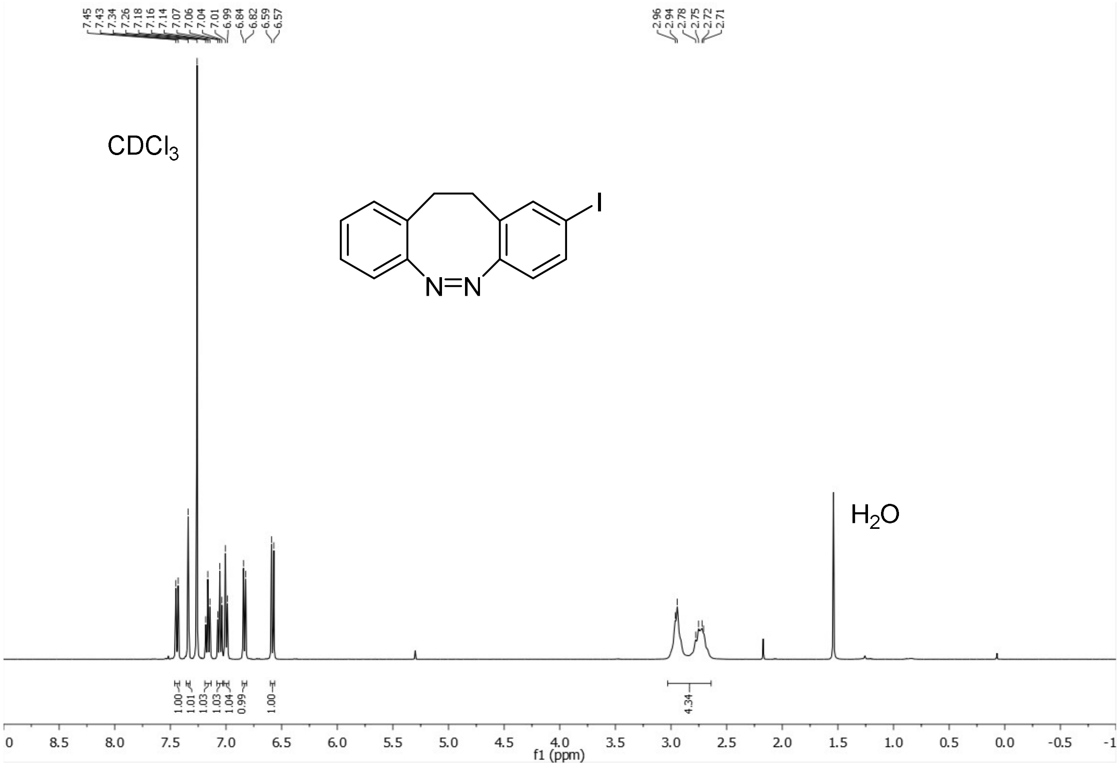


$^{13}$C-NMR (100 MHz, $CDCl_3$) – **3**

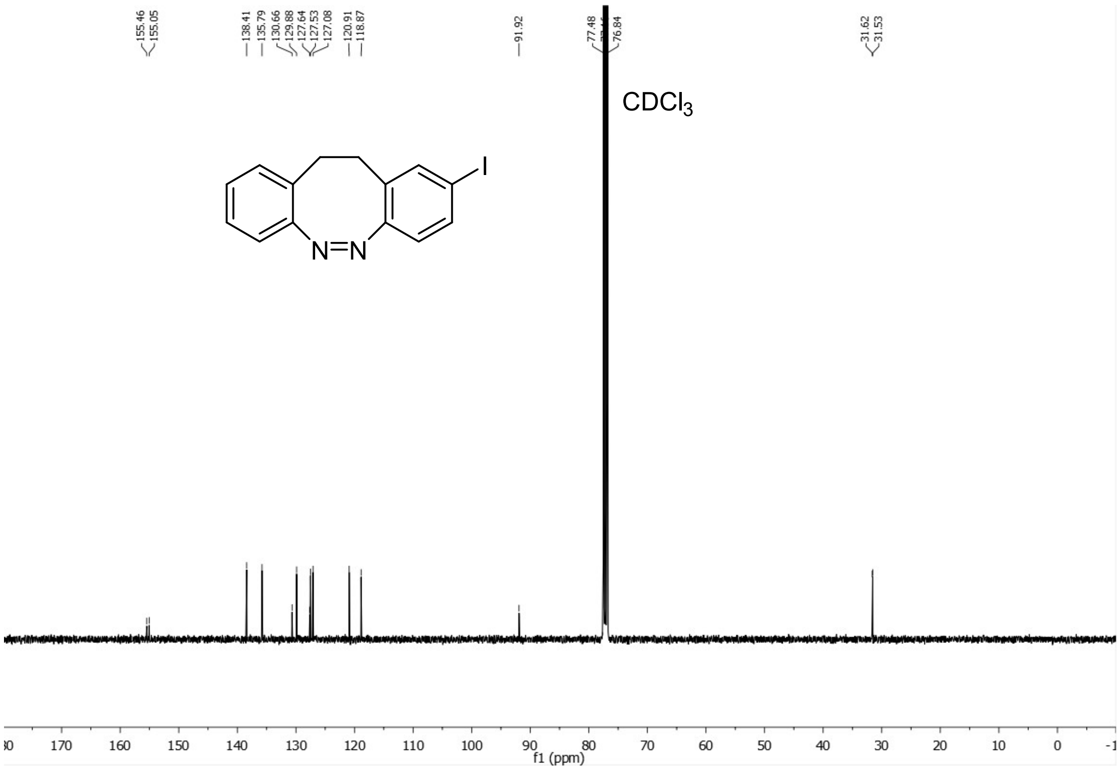

$^1H$-NMR (400 MHz, $CDCl_3$) – **5**

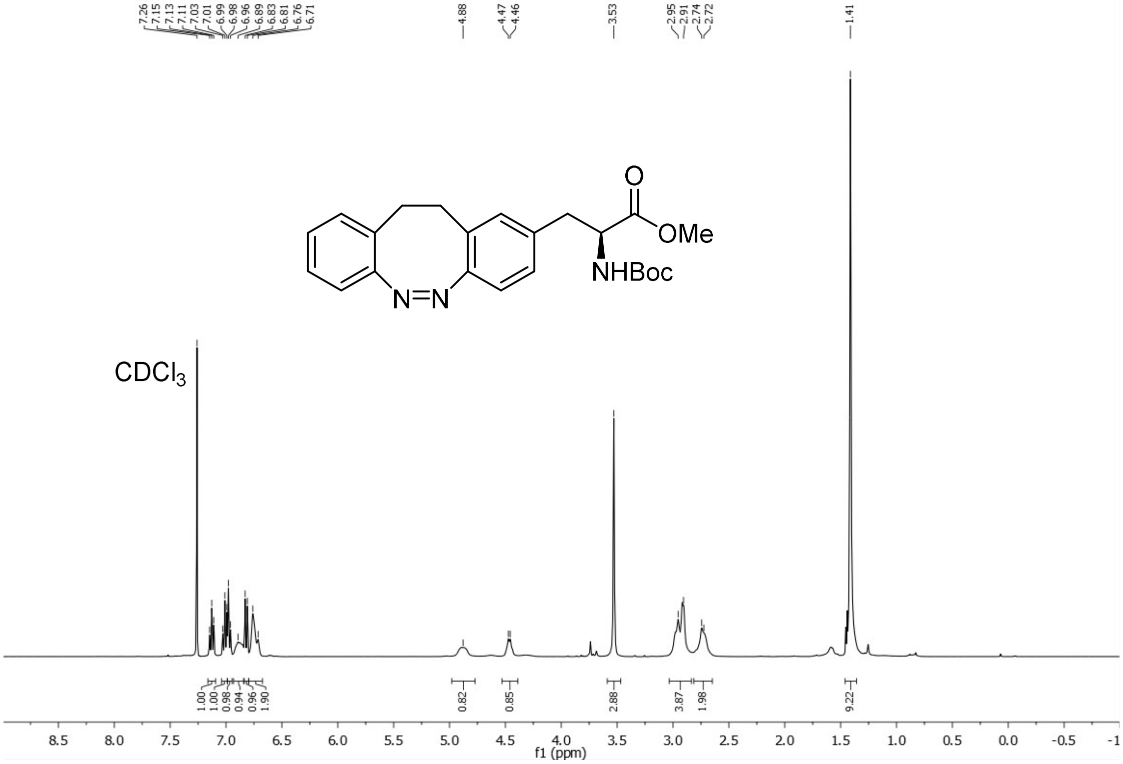


$^{13}C$-NMR (100 MHz, $CDCl_3$) – **5**

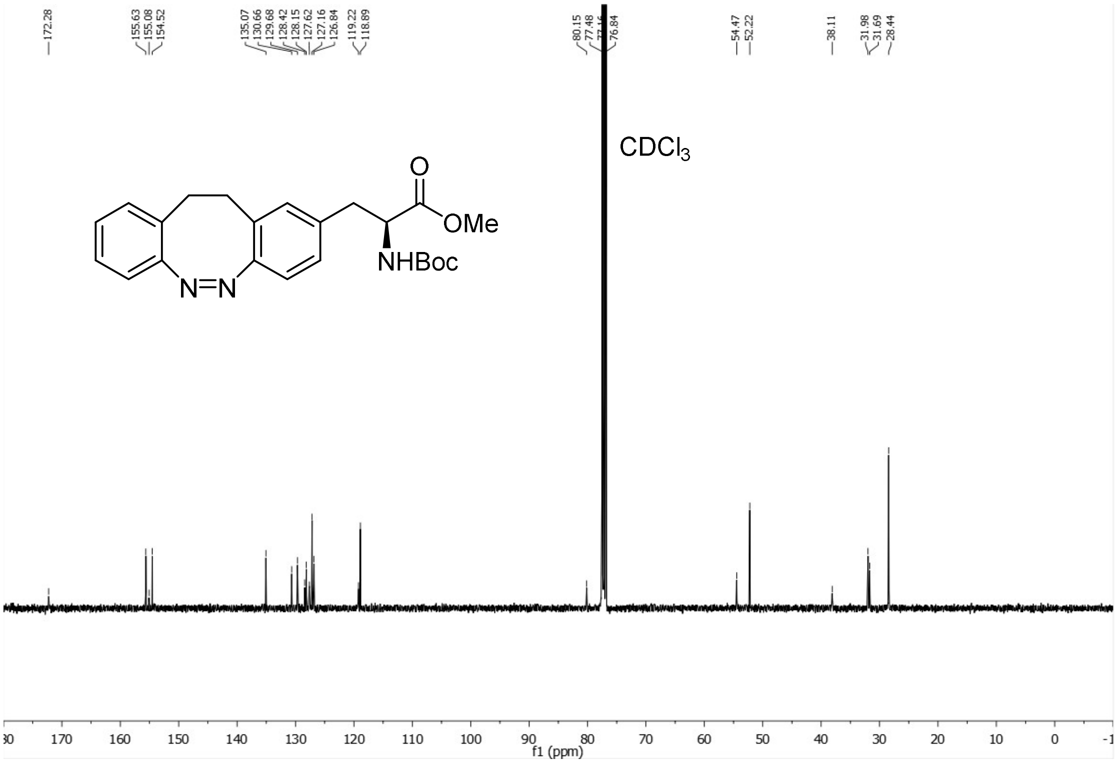

$^{1}$H-NMR (400 MHz, $CDCl_3$) – **S4**

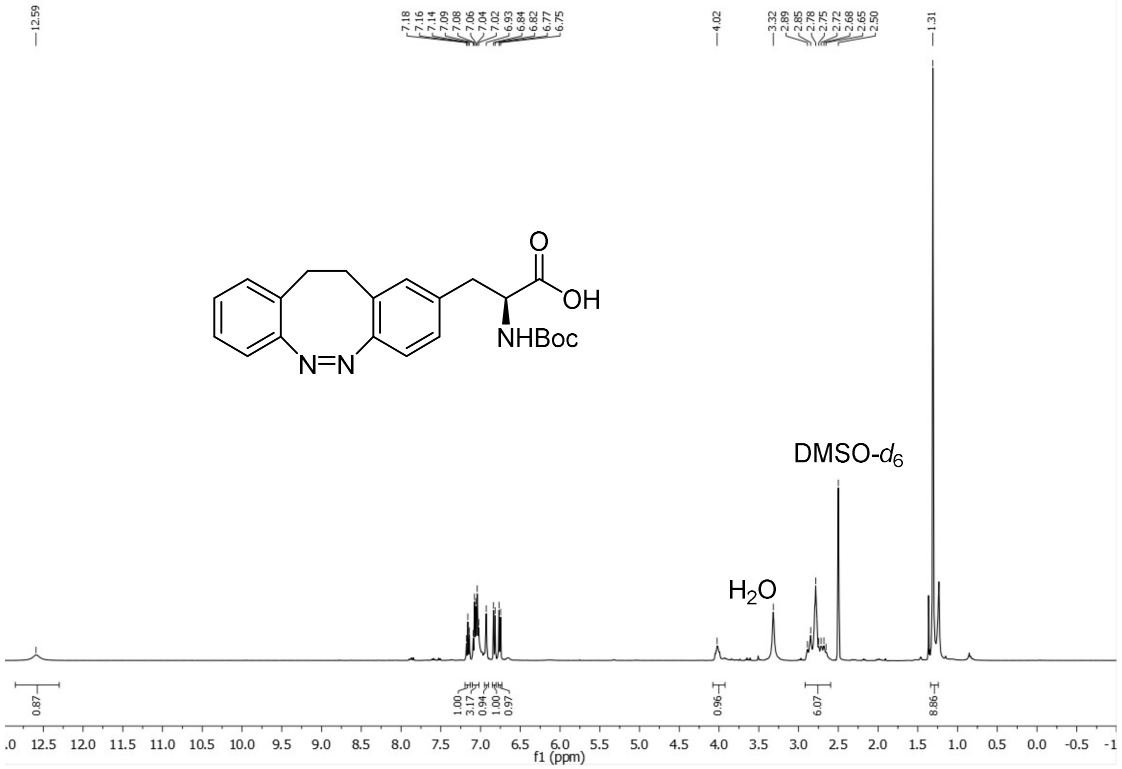


$^{13}$C-NMR (100 MHz, $CDCl_3$) – **S4**

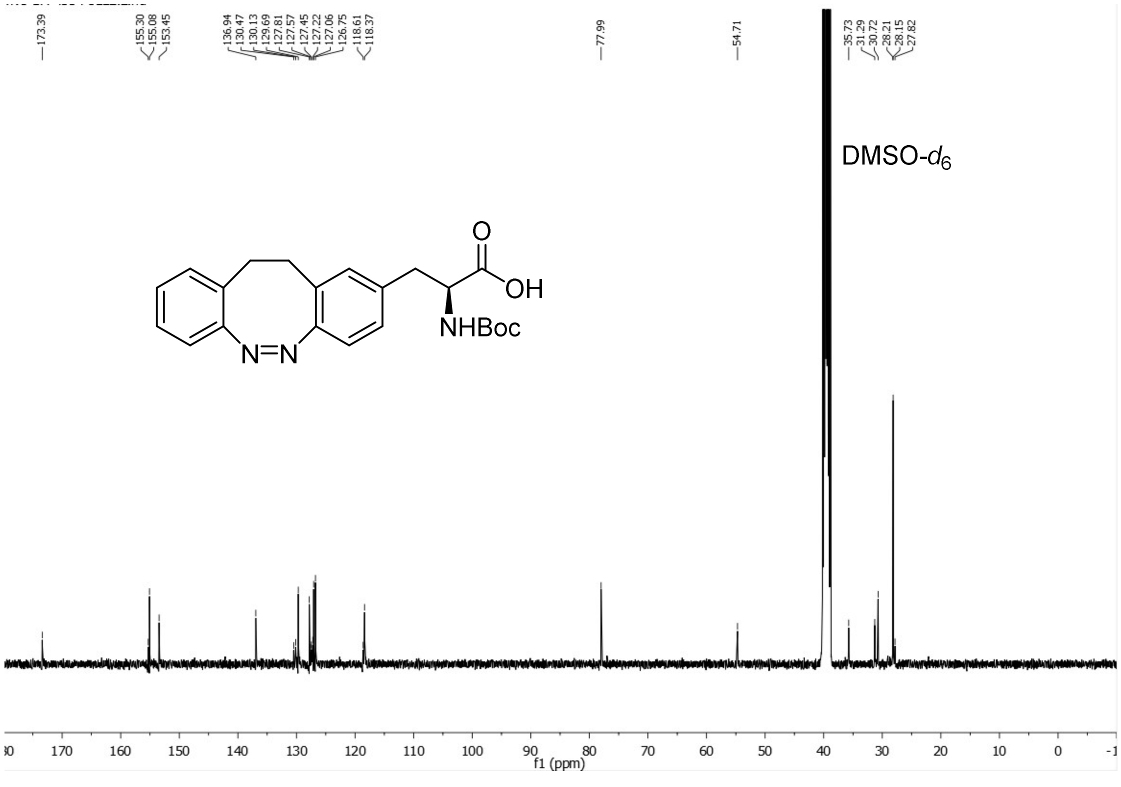

[1]H-NMR (400 MHz, $CDCl_3$) – **6**

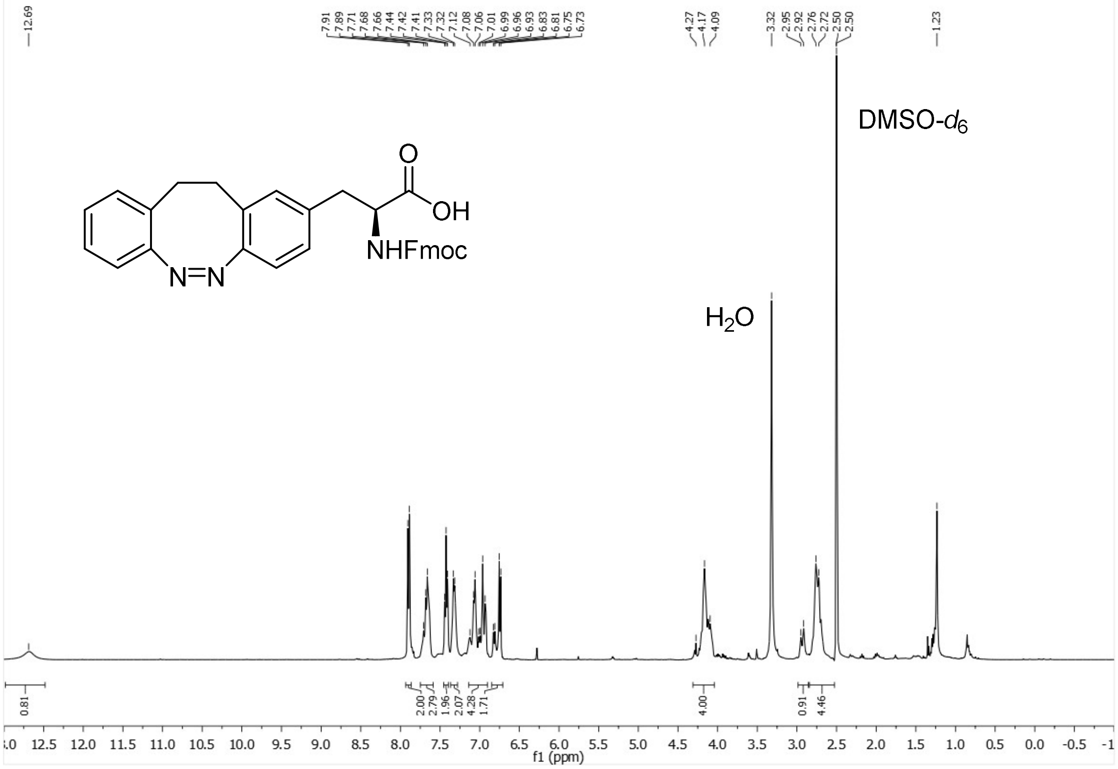


[13]C-NMR (100 MHz, $CDCl_3$) – **6**

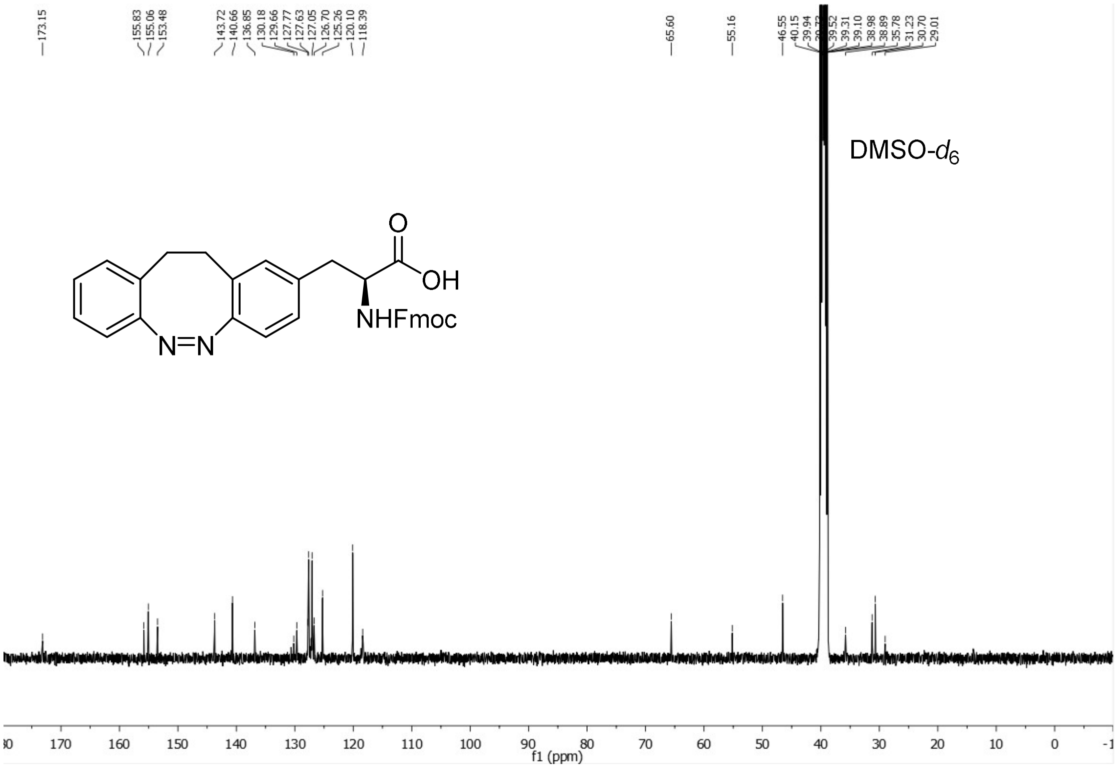

$^1$H-NMR (400 MHz, $CDCl_3$) – **7**

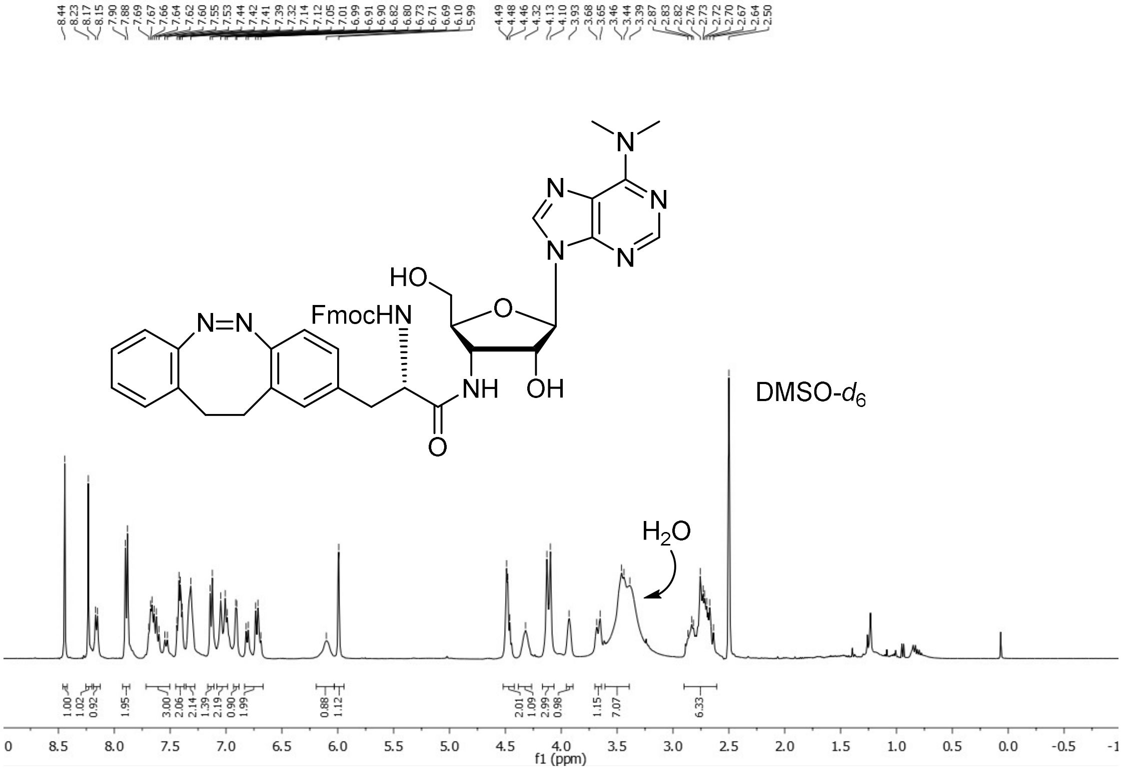


$^{13}$C-NMR (100 MHz, $CDCl_3$) – **7**

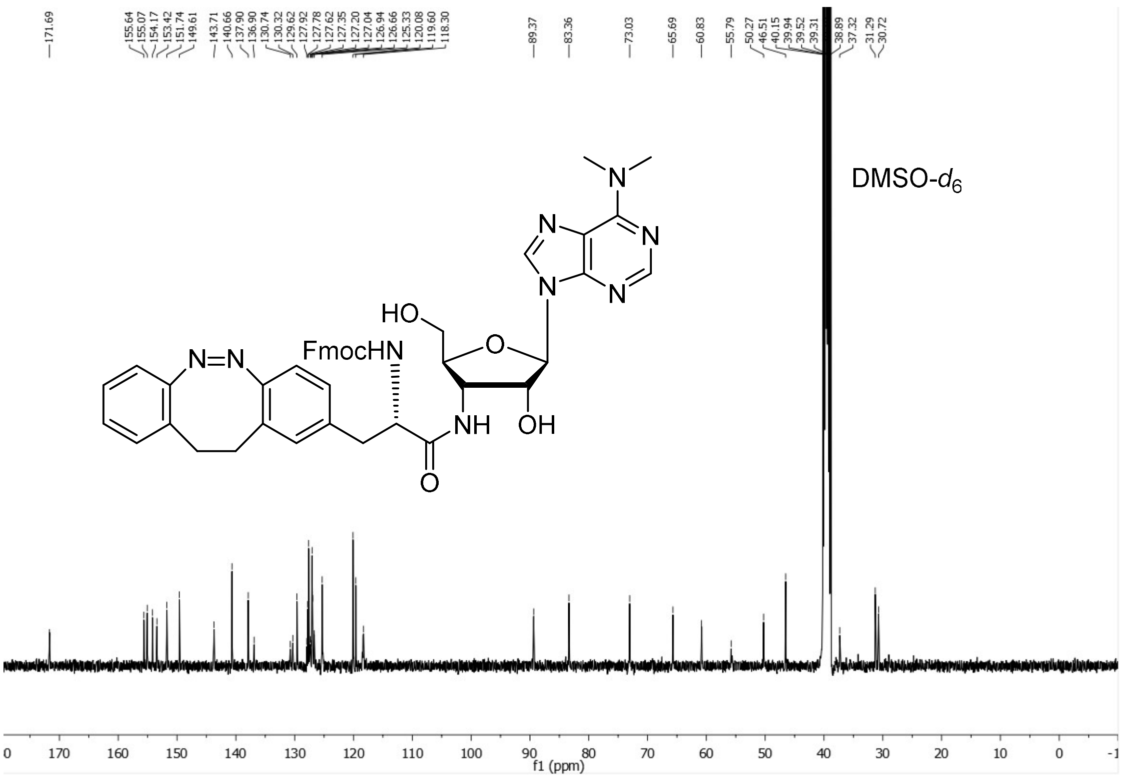


$^{1}$H-NMR (400 MHz, $CDCl_3$) – **puroswitch**

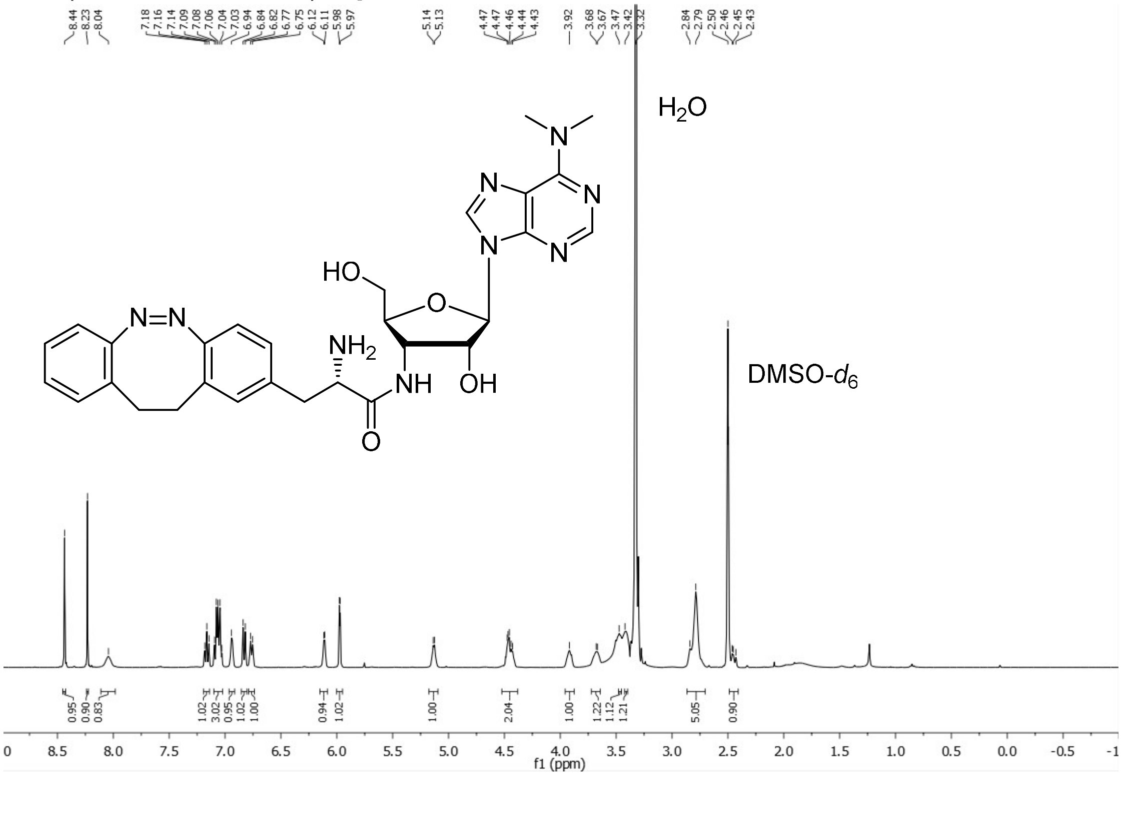


$^{13}$C-NMR (100 MHz, $CDCl_3$) – **puroswitch**

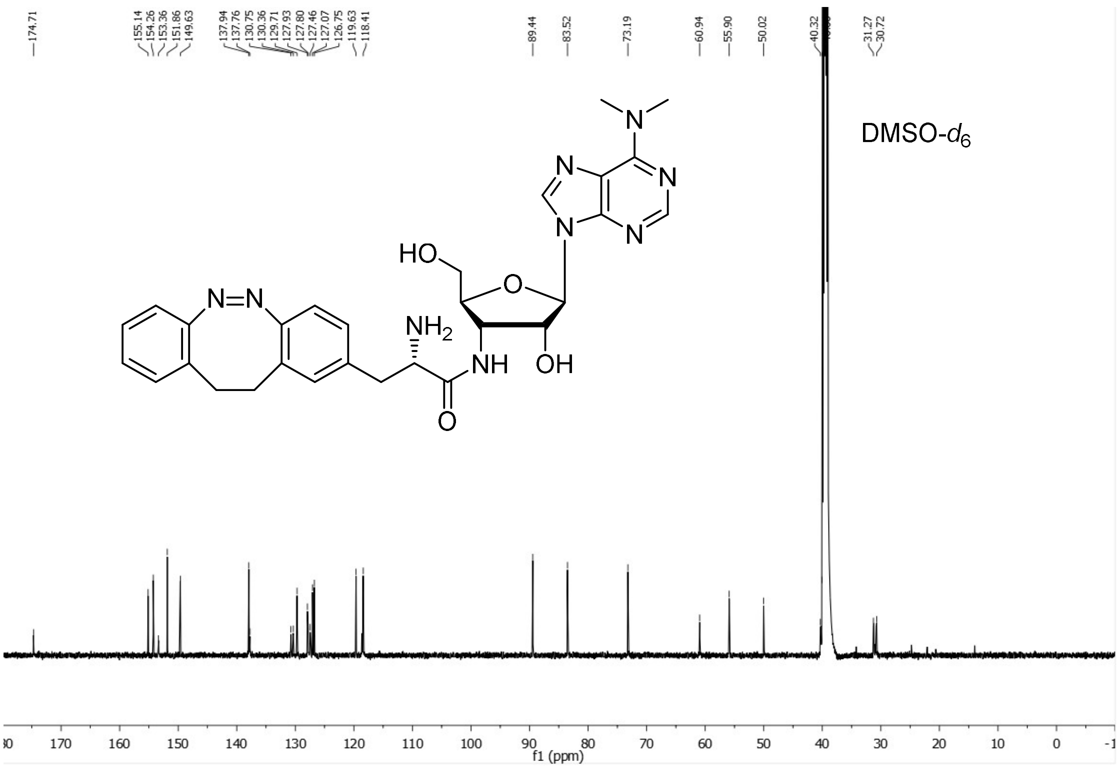

174.71
155.14
154.26
153.36
151.86
149.63
137.94
137.76
130.75
130.36
129.71
127.93
127.80
127.46
127.07
126.75
119.63
118.41
89.44
83.52
73.19
60.94
55.90
50.02
40.32
31.27
30.72
DMSO-$d_6$
f1 (ppm)
